# Population productivity of wedgefishes, guitarfishes, and banjo rays: inferring the potential for recovery

**DOI:** 10.1101/584557

**Authors:** Brooke M. D’Alberto, John K. Carlson, Sebastián A. Pardo, Colin A. Simpfendorfer

## Abstract

Recent evidence of widespread and rapid declines of wedgefishes, guitarfishes, and banjo ray populations (Order Rhinopristiformes), driven by a high demand for their fins in Asian markets and the quality of their flesh, raises concern about their risk of over-exploitation and extinction. Using life history theory and incorporating uncertainty into a modified Euler-Lotka model, maximum intrinsic rates of population increase (*r*_*max*_) were estimated for nine species from the four families of rhinopristiforms. Estimates of median *r*_*max*_ varied from −0.04 to 0.60 year^−1^ among the nine species, but generally increased with increasing maximum size. In comparison to 115 other species of chondrichthyans for which *r*_*max*_ values were available, the families Rhinidae and Glaucostegidae are relatively productive, while most species from Rhinobatidae and Trygonorrhinidae had relatively low *r*_*max*_ values. If the demand for their high value products can be addressed, then population recovery for this species is likely possible but will vary depending on the species.

## Introduction

Understanding the ability of species to recover from declines following implementation of management measures is important to rebuilding depleted populations. This can be approximated through measuring the species’ population productivity using various demographic techniques such as rebound potential models [1–3], age or stage structured life history tables and matrix models [4, 5], and demographic invariant methods [6, 7]. These demographic techniques utilise the known relationships between life history traits and demography, known as the Beverton-Holt dimensionless ratios [8] that can be used to infer a species’ ability to rebound following population declines [9–11]. One commonly used parameter is the maximum intrinsic rate of population increase *r*_*max*_, which reflects the theoretical productivity of depleted populations in the absence of density dependent regulation [12]. This method can be used to provide insights into demography and fisheries sustainability [13], particularly for poorly monitored species with limited available information, like many chondrichthyan (sharks, rays, and chimeras, Class Chondrichthyes) populations [14, 15].

An estimated 25% of chondrichthyans populations have an elevated risk of extinction [16], raising significant ecological and conservation concerns [17–19]. Chondrichthyans, generally, have low biological productivity (slow growth, late maturity, few offspring, and long generational times), which limits their recovery from population declines [20, 21]. Declines of chondrichthyan populations are typically the result of the rapid expansion of fisheries [22–24] and the globalisation of trade [25, 26], and can be exacerbated by habitat degredation [27]. Among species, larger elasmobranchs (sharks and rays, Subclass Elasmobranchii) have some of the lowest intrinsic rates of population increase [28, 29], and as a result are unlikely to sustain high levels of fishing pressure before population collapse [30–33].

The order Rhinopristiformes, is considered one of the most threatened orders of chondrichthyans [16]. All five species of sawfishes (Pristidae), five out of six species of giant guitarfishes (Glaucostegidae), seven out of 10 species of wedgefishes (Rhinidae), six of 31 species of guitarfishes (Rhinobatidae) and two of eight species of banjo rays (Trygonorrhinidae) are listed in a Threatened category (i.e. Vulnerable, Endangered, or Critically Endangered) on the International Union for Conservation of Nature’s (IUCN) Red List of Threatened Species [34]. These large rays are strongly associated with soft-bottom habitats in shallow (<100 m) tropical and temperate coastal waters [35–37], resulting in high exposure to intensive and expanding fisheries, which can also result in habitat degradation [38, 39]. These groups are extremely sensitive to overexploitation as a result of their large body size, slow life history [16] and use of inshore habitat in some of the world’s most heavily fished coastal regions [40–42]. There is increasing evidence of extensive and rapid declines in populations of wedgefishes and giant guitarfishes throughout most of their ranges, including Indonesia [43], South Africa [44], Madagascar [45], Mozambique [46], Tanzania [47] Arabian Seas and surrounding region [34, 48], India [49] and Brazil [50].

The widespread declines of wedgefishes and giant guitarfishes are linked to the international shark fin trade, as their fins are considered amongst the most lucrative shark and ray products [22, 38, 43, 51]. The fins of wedgefishes and giant guitarfishes are considered the highest grade fins and are prevalent in shark fin trading hubs such as Hong Kong [52] and Singapore [53, 54]. Given high demand of the shark fin trade and increasing evidence of population declines, there is considerable concern that wedgefishes and giant guitarfishes are following a similar pattern of global decline as the sawfishes [34]. All five species of sawfish declined rapidly over 30 years throughout their range, driven by unregulated fisheries, the interational fin trade, and delayed scientific attention [55–58]. Yet despite a global conservation strategy [38], restriction of international trade (i.e. listing on CITES Appendix I), and evidence that some species of sawfish have the ability to recover from fishing pressure in the foreseeable future [59], the recovery of the populations is projected to take at least several decades. To avoid the same fate of sawfishes, precautionary management and conservation for wedgefishes, guitarfishes, and banjo rays will be vital to maintain populations.

Currently, wedgefishes, guitarfishes (including families Glaucostegidae and Rhinobatidae), and banjo ray fisheries and trade are not regulated through species-specific fishing or trade restrictions. The magnitude of the declines and the subsequent conservation issues have attracted the focus of major international management conventions and agencies, with *Rhynchobatus australiae* and *Rhinobatos rhinobatos* listed on the Appendix II of Convention on the Conservation of Migratory Species of Wild Animals (CMS) in 2017 [60]. In addition, *R. australiae, Rhynchobatus djiddensis, Rhynchobatus laevis*, and *R. rhinobatos* were listed on Annex 1 of the CMS Memorandum of Understanding (MOU) on the Conservation of Migratory Sharks in 2018 [61], and the families Rhinidae and Glaucostegidae have been proposed for listing on the Appendix II of the Convention on International Trade in Endangered Species (CITES) [62]. Given the global concerns for this group of species, and the importance of trade in their high value fins, the use of trade regulations through CITES listings may help achieve positive conservation outcomes [63]. However, management and
conservation efforts can be hampered by the lack of life history (e.g. age, growth and maturity) and demographic information. Despite the high exposure to fisheries, there is limited information available globally on biology (age, growth, maturity, demography), and abundance and distribution of wedgefishes, guitarfishes, and banjo rays.

Estimating *r*_*max*_ values can provide the demographic basis for evaluating the sustainability of wedgefish and guitarfish fisheries and international trade, and their ability to recover from population declines, while accounting for uncertainty about their life histories [28]. While the maximum intrinsic population rate of population increase has previously been estimated for *Pseudobatos horkelii* and *Pseudobatos productus* as a part of multi-species comparison [15, 64], there has not been a comprehensive analysis on the population productivity for rhinopristiforms.The focal wedgefishes, guitarfishes, and banjo ray families studied in this paper were Rhinidae (wedgefishes), Glaucostegidae (giant guitarfishes), Rhinobatidae (guitarfishes) and Trygonorrhinidae (banjo rays), while the family Pristidae (sawfishes) were excluded as they have been previously assessed in detail by. Dulvy et al. (58). The aim of this paper was to use life history data and theory, and comparative analysis, to estimate the population productivity for wedgefishes, guitarfishes, and banjo rays, particularly within the context of other sharks and rays.

## Materials and Methods

### Life history data collection

A literature search was conducted for all species from four families of Rhinopristiformes: Rhinidae, Glaucostegidae, Rhinobatidae and Trygonorrhinidae, to provide data for estimation of population productivity. Life history information required for analyses consisted of age at maturity (*α*_*mat*_, range of years), maximum age (*α*_*max*_, in years), range of litter size (in number of female pups), sex ratio, breeding intervals (*i*, years), and von Bertalanffy growth coefficient (*k*, year^−1^). Out of the four families, with a total of 57 species, only nine species had enough published life history information to estimate *r*_*max*_ (Table 1).

**Table 1.**
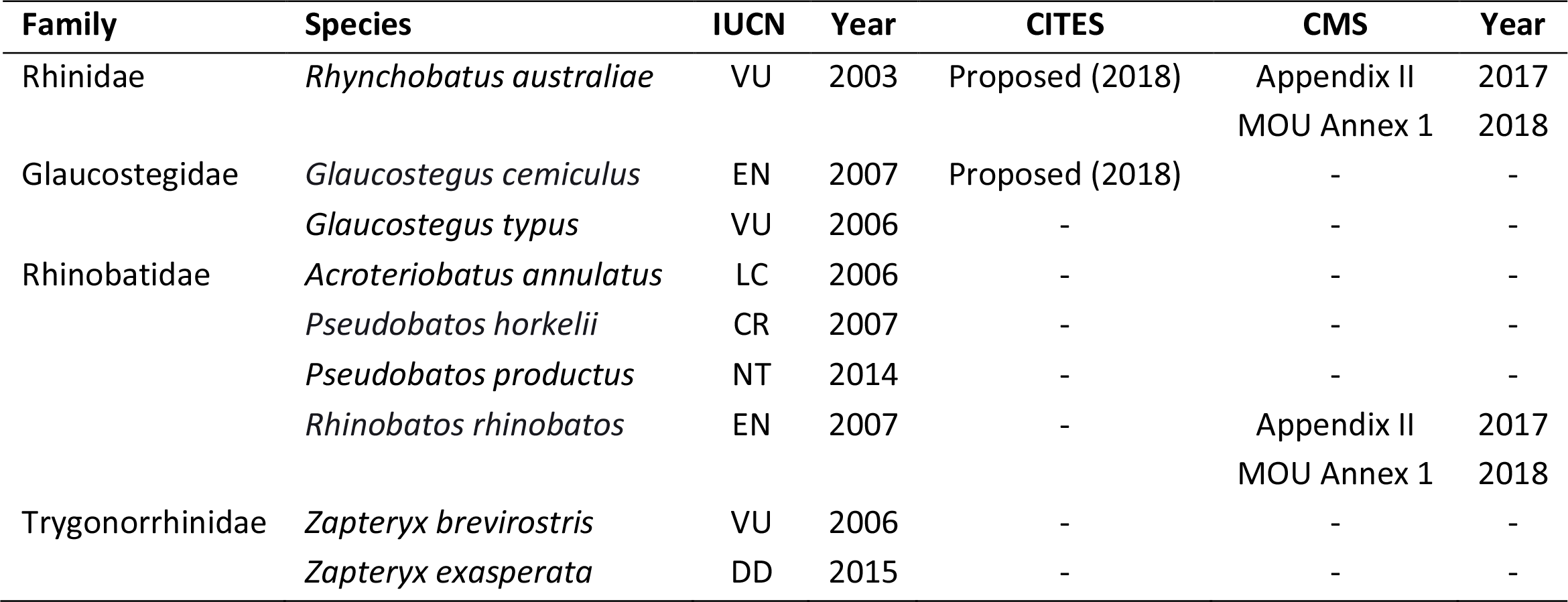
The nine species of wedgefishes, guitarfishes, and banjo rays included in this study, their threat status according the International Union of Conservation of Nature (IUCN)’s Red List of Threatened Species, and whether the species are listed on the appendixes of Convention of International Trade of Endangered Species (CITES) and/or Convention on the Conservation of Migratory Species of Wild Animals (CMS) and the CMS Memorandum of Understanding (MOU). IUCN categories are CR, Critically Endangered; EN, Endangered; VU, Vulnerable; LC, Least Concern; DD, Data Deficient.

Growth coefficient data for *R. australiae* was reported as *Rhynchobatus spp*. by White et al. (65)
as results from the species complex including *R. australiae, Rhynchobatus palpebratus* and *Rhynchobatus laevis* along the eastern coast of Australia. Recent taxonomic revision has resolved this species complex in this area, with *R. laevis* primarily found in the Indian Ocean and Indo-West Pacific Ocean [66], and further examination of data associated with specimen examined by White et al.(65) have demonstrated they were primarily *R. australiae*.

For *R. australiae*, *G. typus* and *Z. brevirostris* the age at maturity was back-calculated using:

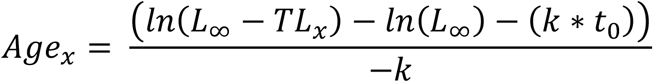

where *Age*_*x*_ is age at time x, *TL*_*x*_ is total length (cm TL) at time x, *L*_∞_ is the asymptotic length (cm TL), *t*_0_ is the length at time zero, and *k* is the von Bertalanffy growth coefficient. For *R. australiae*, the age at maturity was back-calculated using the von Bertalanffy parameters reported for *Rhynchobatus spp*. by White et al. (65) and the size at maturity of 150cm TL from *Rhynchobatus djiddensis* [66]. The age at maturity for *Glaucostegus typus* was estimated using the estimated size at maturity in Last et al. (66) and growth coefficient in White et al. (65). There is no reported litter size for *G. typus*, thus we assumed to have the same litter size and breeding interval as *Glaucostegus cemiculus* to calculate annual reproductive output. For *R. australiae*, *Acroteriobatus annulatus, Zapteryx exasperata* and *Z. brevirostris*, the breeding interval was assumed to be one year, as there was no information available (Table 2).

**Table 2.** Life history values and sources used to estimate maximum intrinsic rate of population increase (*r*_*max*_) for the nine species of wedgefishes, guitarfishes, and banjo rays studied, including the maximum size (Max. size in centimetres total length, cm TL), lower, upper and mean (standard deviation, S.D.) values of the age at maturity (*α*_*mat*_, years), lower and upper values for litter size, breeding interval (*i*, years), lower and upper annual reproductive output of females (*b*), lower and upper values for von Bertalanffy growth coefficient (*k*,year^−1^), the observed, and lower (*T*_*lower*_) and upper (*T*_*upper*_) and mean (S.D.) values of theoretical maximum age (*α*_*max*_, years).

| Family | Species | Max.<br>size | $\alpha_{mat}$ | | | | litter size | | i | $b$ | | $k$ | | $\alpha_{max}$ | | | | | References |
| --- | --- | --- | --- | --- | --- | --- | --- | --- | --- | --- | --- | --- | --- | --- | --- | --- | --- | --- | --- |
| | | (cm<br>TL) | lower | upper | mean | ± S.D. | lower | upper | (years) | lower | upper | lower | upper | Observed | $T_{lower}$ | $T_{upper}$ | mean | ± S.D. | |
| Rhinidae | *Rhynchobatus australiae* | 300 | 3.00 | 6.00 | 4.50 | 0.450 | 7 | 19 | 1 | 3.5 | 9.5 | 0.400 | 0.400 | 12.0 | 11.3 | 11.3 | 11.51 | 0.346 | [65, 66] |
| Glaucostegidae | *Glaucostegus cemiculus* | 290 | 2.89 | 6.50 | 4.70 | 0.680 | 5 | 24 | 1 | 2.5 | 12 | 0.200 | 0.275 | 14.0 | 13.9 | 16.1 | 14.67 | 0.50 | [66-70] |
|  | *Glaucostegus typus* | 270 | 6.50 | 8.00 | 7.25 | 0.245 | 5 | 24 | 1 | 2.5 | 12 | 0.150 | 0.150 | 19.0 | 18.1 | 18.1 | 18.42 | 0.16 | [65, 66, 71] |
| Rhinobatidae | *Acroteriobatus annulatus* | 140 | 2.30 | 2.80 | 2.55 | 0.080 | 2 | 10 | 1 | 1.0 | 5.0 | 0.240 | 0.240 | 7.00 | 14.8 | 14.8 | 12.23 | 1.30 | [66, 72] |
|  | *Pseudobatos horkelii* | 140 | 7.00 | 9.00 | 8.00 | 0.300 | 4 | 12 | 1 | 2.0 | 6.0 | 0.194 | 0.194 | 28.0 | 16.3 | 16.3 | 22.17 | 1.86 | [66, 73] |
|  | *Pseudobatos productus* | 170 | 7.00 | 8.40 | 7.70 | 0.200 | 1 | 10 | 1 | 0.5 | 5.0 | 0.016 | 0.240 | 33.8 | 14.8 | 33.8 | 33.80 | 3.50 | [66, 74-76] |
|  | *Rhinobatos rhinobatos* | 185 | 2.20 | 4.10 | 3.15 | 0.350 | 1 | 14 | 1 | 0.5 | 7.0 | 0.134 | 0.310 | 18.9 | 13.1 | 18.9 | 18.92 | 1.00 | [66, 77-82] |
| Trygonorrhinidae | *Zapteryx brevirostris* | 66.0 | 7.71 | 11.5 | 9.61 | 0.700 | 1 | 8 | 1 | 0.5 | 4.0 | 0.110 | 0.130 | 10.0 | 19.1 | 20.3 | 16.48 | 1.55 | [66, 83-86] |
|  | *Zapteryx exasperata* | 103 | 5.41 | 9.65 | 7.53 | 0.800 | 2 | 13 | 1 | 1.0 | 6.5 | 0.144 | 0.174 | 22.6 | 17.1 | 18.4 | 19.85 | 0.80 | [66, 87-90] |

### Estimation of maximum intrinsic population growth rate, *r*_*max*_

Maximum intrinsic rate of population increase was estimated using an unstructured derivation of the Euler-Lotka model. This model accounts for juvenile survivorship that depends on age at maturity and species-specific natural mortality, and incorporates uncertainty within the parameters through Monte Carlo simulation [64, 91]. Requirements of this model are estimates of three biological parameters: annual reproductive output, age at maturity, and natural mortality. This model is founded on the principle that a breeding female only has to produce one mature female in her lifetime to ensure a stable population [92–95]: 
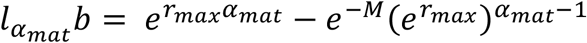

where 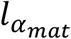 is survival to maturity in the absence of fishing and is calculated as 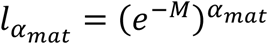, b is the annual rate of production of females, *α*_*mat*_ is the age of maturity and *M* is instantaneous natural mortality. The annual reproductive output of females was calculated as *b*= 0.5*l*/*i*, where *l* is litter size (in number of males and females) and *i* is breeding interval (in years). Annual reproductive output estimates were derived from uniform distributions constrained by the minimum and maximum litter sizes published in the literature (Table 2). If the litter sex ratio was unknown, it was assumed to be 1:1. Age at maturity estimates were derived from normal distributions with means and standard deviations (S.D.) calculated from the available ages at maturity published in the literature for each species (Table 2). Normal distributions were truncated to be positive and within “reasonable bounds”. The von Bertalanffy growth coefficients (*k*) for each species were derived from uniform distributions ranging between the minimum and maximum published values (Table 2). As the observed maximum age may not reflect the longevity of the species [96], the theoretical maximum age (*T*_*max*_) was calculating using minimum and maximum *k* reported for each species in the literature, using the following the formula from Mollet et al. (97):

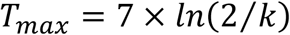

Maximum age (*α*_*max*_) estimates were derived from a normal distribution using the mean and S.D., calculated from the observed maximum age reported in the literature, minimum theoretical maximum age (*T*_*lower*_) and maximum theoretical age (*T*_*upper*_). As there was no current consensus on the best indirect method to estimate the instantaneous natural mortality, it was estimated using four common methods, Jensen’s First mortality estimate [98], modified Hewitt and Hoeing estimator [99], Frisk’s estimator [9], and reciprocal of the lifespan [10] (Table 3).

**Table 3.** Natural mortality (*M*) methods used to estimate maximum intrinsic rate of population increase (*r*_*max*_) in this study, where *α*_*mat*_ is age at maturity in years, *α*_*max*_ is maximum age in years, and *k* is the von Bertalanffy growth coefficient in year^−1^.

| Method | Equation | References |
| --- | --- | --- |
| 1. Jensen's First Estimator | $M = 1.65/\alpha_{mat}$ | Jensen (98) |
| 2. Modified Hewitt & Hoeing Estimator | $M = 4.22/\alpha_{max}$ | Hewitt and Hoenig (99) |
| 3. Frisk's Estimator | $M = 0.4/k$ | Frisk et al. (9) |
| 4. Reciprocal of lifespan | $M = 1/(\alpha_{mat} + \alpha_{max}/2)$ | Pardo et al. (64) |

The annual reproductive output and age at maturity were highly uncertain parameters, while the natural mortality was estimated indirectly, which can result in additional uncertainty [28]. Monte Carlo simulation was used to account for uncertainty of input parameters. Model parameters were drawn from their respective distributions iteratively 20,000 times [14]. To incorporate uncertainty into *M*, for each iteration the values for age at maturity, maximum age and von Bertalanffy growth coefficients were drawn from their respective distributions and used to estimate natural mortality for the four natural mortality estimators, which in turn is required to estimate *r*_*max*_ [14]. Scenarios were investigated where uncertainty was only incorporated into a single parameter. Values of one parameter were drawn from its distribution, while the remaining parameters were set as deterministic by using the median values of their respective distributions. This was done for the age at maturity, annual reproductive output and natural mortality. The *M* value was set as a deterministic in the other scenarios, even when the parameters used to estimate *M* were being drawn from distributions. In each iteration, the *r*_*max*_ equation was solved using the *nlminb* optimisation function by minimising the sum of squared differences. This range of *r*_*max*_ values was generated to encompass the widest range of plausible life histories and should therefore include the true parameter values. This method was applied to each of the nine species for which data were available and from this range, median and mean *r*_*max*_ values and standard deviation were calculated.

### Comparison of wedgefishes, guitarfishes, and banjo ray *r*_*max*_ estimates among chondrichthyans

Median *r*_*max*_ of the nine wedgefishes, guitarfishes, and banjo ray species were compared to all available estimates using recalculated values by Pardo et al. (64) to incorporate survival to maturity, including an additional 14 species (S1 Table). Following the described method above, the median *r*_*max*_ was calculated for the additional species, including great hammerhead *Sphyrna mokarran*, smooth hammerhead *Sphyran zygaena*, common thresher shark *Alopias vulpinus*, reef manta ray *Mobula alfredi*, giant manta ray *Mobula birostris*, Chilean devilray *Mobula tarapacana*, bentfin devil *Mobula thurstoni*, blackspotted whipray *Maculabatis astra*, speckled maskray *Neotrygon picta*, narrow sawfish *Anoxypristis cuspidata*, dwarf sawfish *Pristis clavata*, smalltooh sawfish *Pristis pectinata*, green sawfish *Pristis zijsron*, and Atlantic angel shark *Squantina dumeril* (S1 Table). The reciprocal of the lifespan natural mortality method was chosen to estimate the natural morality in order to compare to values generated by Pardo et al. (64) as that was the method used in their study. The *r*_*max*_ estimates for *Pseudobatos horkelii* and *Pseudobatos productus* were updated with the values from this study for the comparison. The age at maturity (years), maximum age (years), growth rate (k, years^−2^) and maximum size in centimetres (cm) were plotted against the *r*_*max*_ estimates for 115 chondrichthyan species, including the nine species of wedgefishes, guitarfishes, and banjo rays (S1 Table). Maximum sizes were total length (TL) for all species except for Myliobatiformes, where the disc width (DW) were used [15, 28]. All models and figures were built in the R version 3.4.1 version [100] (S1 Table).

## Results

Estimates of maximum intrinsic rate of population increase for the nine species of wedgefishes, guitarfishes, and banjo ray varied considerably among species, between families and by the method of estimating natural mortality. There was a high level of uncertainty in the annual reproductive output and age at maturity across all species (Fig 1). Uncertainty in the natural mortality values were low (Fig 1), but it resulted in high uncertainty in the *r*_*max*_ estimates, which was highly influenced by the natural mortality estimator (Fig 2; Table 4).

**Figure 1.**
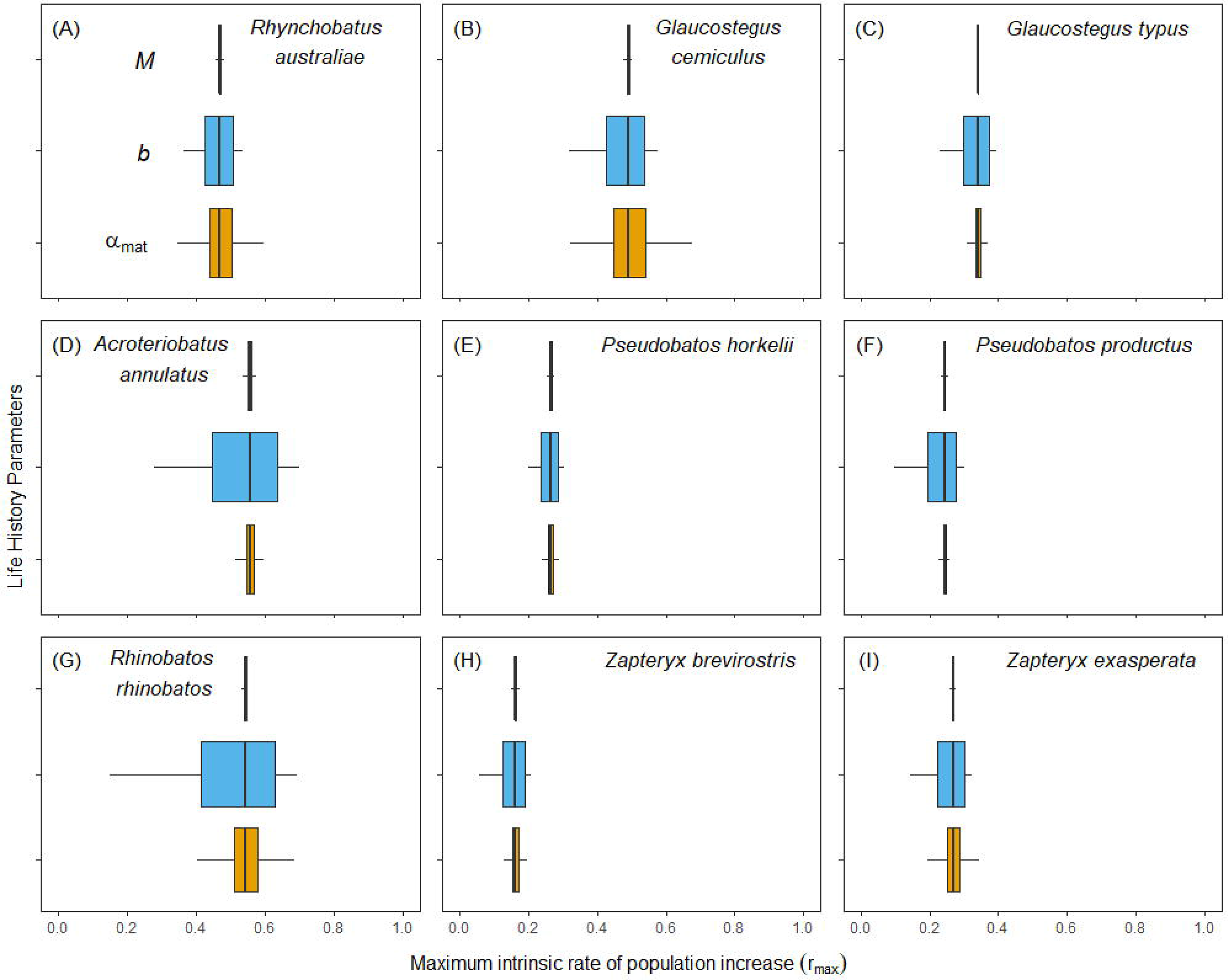
Predicted values of maximum intrinsic rate of population increase (*r*_*max*_, year^−1^) for nine wedgefishes, guitarfishes, and banjo ray species when including uncertainty in the reciprocal of the lifespan natural mortality estimator (*M*, top/grey boxplot), annual reproductive output (*b*, middle/blue boxplot), and age at maturity (*α*_*mat*_, bottom/orange boxplot). Specie are (A) *Rhynchobatus australiae*, (B), *Glaucostegus cemiculus*, (C) *Glaucostegus typus*, (D) *Acroteriobatus annulatus*, (E) *Pseudobatos horkelii*, (F) *Pseudobatos productus*, (G) *Rhinobatos rhinobatos*, (H) *Zapteryx brevirostris*, and (I) *Zapteryx exasperata*. The natural mortality estimator used for this graph is the. Boxes indicate median, 25 and 75% quantiles, whereas the lines encompass 95% of the values (2.5 and 97.5% quantiles). For plots incorporating uncertainty with other three natural mortality methods, see Supplementary Material, S2-S4 Figures.

The ranges of *r*_*max*_ for each species were relatively large as a result of the high uncertainty in the life history parameters and method of estimating natural mortality (Fig 2). *Acroteriobatus annulatus* and *R. rhinobatos* had the largest range of *r*_*max*_, regardless of the natural mortality estimation method used (Fig 2; Table 4). *Pseudobatos horkelii* and *P. productus* had the smallest range of *r*_*max*_ (Fig 2; Table 4). For all species, the highest *r*_*max*_ values were estimated using the maximum age natural mortality method, excluding *P. productus* and *Z. brevirostris*, which had highest *r*_*max*_ values estimated from Frisk’s Estimator (Table 4). Frisk’s estimator, Maximum Age and Lifespan methods produced similar *r*_*max*_ estimates for each species, with 7% or less difference between median values (Table 4, Fig 2).

**Table 4.** Maximum intrinsic rates of population increase estimates (*r*_*max*_, year^−1^) for nine species of wedgefishes, guitarfishes, and banjo ray, using four estimators of natural mortality. The minimum (Min.), median, mean (± S.E.), and maximum (Max.) *r*_*max*_ values are reported for each species and natural mortality estimator.

| Family | Species | Jensen's First estimator |  |  |  |  | Hewitt & Hoeing's estimator |  |  |  |  | Frisk's estimator |  |  |  |  | Reciprocal of lifespan |  |  |  |  |
| --- | --- | --- | --- | --- | --- | --- | --- | --- | --- | --- | --- | --- | --- | --- | --- | --- | --- | --- | --- | --- | --- |
| | | Min. | Median | Mean | ±S.E. | Max. | Min. | Median | Mean | ±S.E. | Max. | Min. | Median | Mean | ±S.E. | Max. | Min. | Median | Mean | ±S.E. | Max. |
| Rhinidae | *Rhynchobatus australiae* | 0.12 | 0.23 | 0.22 | 0.0004 | 0.35 | 0.02 | 0.22 | 0.22 | 0.0005 | 0.59 | 0.24 | 0.43 | 0.43 | 0.0005 | 0.79 | 0.29 | 0.47 | 0.47 | 0.0005 | 0.81 |
| Glaucostegidae | *Glaucostegus cemiculus* | 0.02 | 0.24 | 0.23 | 0.0005 | 0.48 | 0.04 | 0.30 | 0.30 | 0.0005 | 0.86 | 0.23 | 0.49 | 0.49 | 0.0007 | 1.00 | 0.24 | 0.48 | 0.49 | 0.0007 | 1.00 |
|  | *Glaucostegus typus* | 0.08 | 0.19 | 0.18 | 0.0003 | 0.26 | 0.06 | 0.19 | 0.18 | 0.0003 | 0.28 | 0.23 | 0.36 | 0.35 | 0.0003 | 0.46 | 0.21 | 0.34 | 0.33 | 0.0003 | 0.43 |
| Rhinobatidae | *Acroteriobatus annulatus* | -0.25 | 0.05 | 0.03 | 0.0008 | 0.20 | -0.14 | 0.30 | 0.28 | 0.0008 | 0.54 | 0.30 | 0.59 | 0.57 | 0.0008 | 0.80 | 0.22 | 0.54 | 0.52 | 0.0008 | 0.75 |
|  | *Pseudobatos horkelii* | 0.06 | 0.13 | 0.12 | 0.0002 | 0.17 | 0.00 | 0.14 | 0.14 | 0.0002 | 0.25 | 0.17 | 0.25 | 0.25 | 0.0002 | 0.34 | 0.18 | 0.26 | 0.26 | 0.0002 | 0.34 |
|  | *Pseudobatos productus* | -0.06 | 0.09 | 0.08 | 0.0004 | 0.15 | -0.06 | 0.13 | 0.12 | 0.0004 | 0.23 | 0.09 | 0.26 | 0.25 | 0.0004 | 0.37 | 0.09 | 0.24 | 0.23 | 0.0004 | 0.33 |
|  | *Rhinobatos rhinobatos* | -0.31 | 0.13 | 0.10 | 0.0010 | 0.32 | -0.04 | 0.38 | 0.35 | 0.0011 | 0.93 | 0.12 | 0.55 | 0.53 | 0.0011 | 1.00 | 0.13 | 0.54 | 0.51 | 0.0011 | 1.00 |
| Trygonorrhinidae | *Zapteryx brevirostris* | -0.05 | 0.07 | 0.06 | 0.0003 | 0.13 | -0.08 | -0.04 | 0.04 | 0.0003 | 0.10 | 0.08 | 0.19 | 0.19 | 0.0003 | 0.31 | 0.04 | 0.16 | 0.16 | 0.0003 | 0.27 |
|  | *Zapteryx exasperata* | -0.05 | 0.12 | 0.11 | 0.0004 | 0.21 | -0.03 | 0.12 | 0.12 | 0.0004 | 0.34 | 0.12 | 0.27 | 0.27 | 0.0004 | 0.49 | 0.12 | 0.27 | 0.26 | 0.0004 | 0.47 |

**Figure 2.**
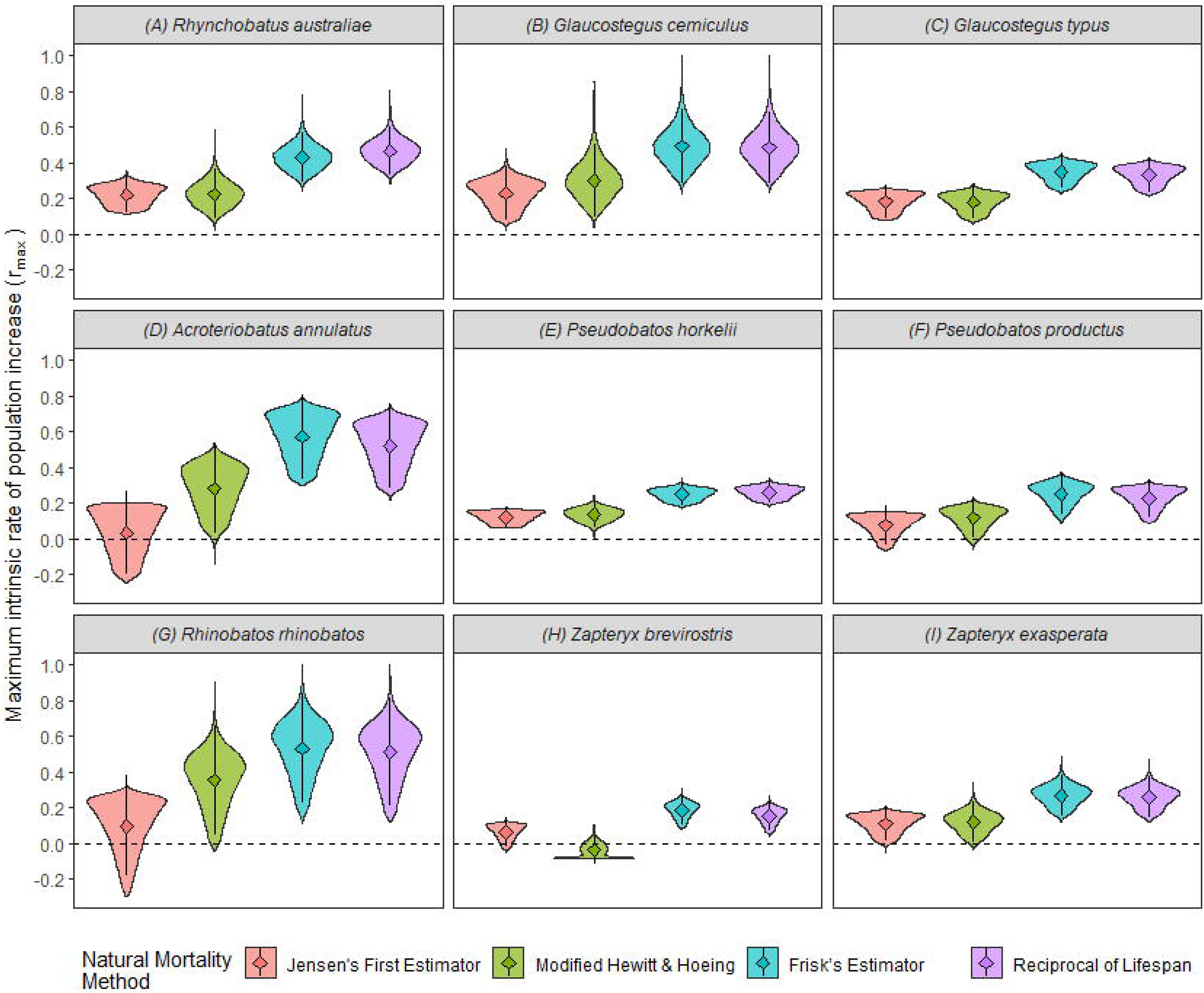
The range of maximum intrinsic rate of population increase (*r*_*max*_, year^−1^) for nine species of wedgefishes, guitarfishes, and banjo rays, obtained with five different methods of estimating the instantaneous natural mortality: Jensen’s First Estimator (red), modified Hoeing & Hewitt’s Estimator (yellow), Frisk’s Estimator (green), and Reciprocal of lifespan (blue). Means (triangle) and standard deviation (black line) are presented for each method. Species are (A) *Rhynchobatus australiae*, (B), *Glaucostegus cemiculus*, (C) *Glaucostegus typus*, (D) *Acroteriobatus annulatus*, (E) *Pseudobatos horkelii*, (F) *Pseudobatos productus*, (G) *Rhinobatos rhinobatos*, (H) *Zapteryx brevirostris*, and (I) *Zapteryx exasperata*. Values below the black dashed line indicate implausible *r*_*max*_ estimates.

*Acroteriobatus annulatus* and *R. rhinobatos* had the highest estimates of *r*_*max*_, followed by *G. cemiculus* and *R. australiae*. *Pseudobatos horkelii, P. productus* and *Z. exasperata* had lower *r*_*max*_ estimates with similar ranges across all natural mortality estimators (Table 4). The lowest *r*_*max*_ values from every species were generated using the Jensen’s First estimator and modified Hewitt and Hoeing’s methods. These methods estimated negative minimum *r*_*max*_ values for *A. annulatus, P. productus, R. rhinobatus, Z. brevirostris* and *Z. exasperata, a*nd a negative median *r*_*max*_ for *Z. brevirostris* (Table 4; Fig 2). *Zapteryx brevirostris*, the smallest species in the study, had one of the lowest estimates of *r*_*max*_, across of natural mortality methods (Table 4).

As the age at maturity decreased, the estimates of maximum intrinsic rate of population increased for the nine species of wedgefishes, guitarfishes, and banjo rays (Fig 3A). The species with the highest median estimates of *r*_*max*_, *R. australiae*, *G. cemiculus*, *R. rhinobatos* and *A. annulatus* had the youngest age at maturity, while *Z. brevirostris* had the oldest age at maturity and lowest median estimate for *r*_*max*_ (Fig 3A). The estimates of *r*_*max*_ increased with the increasing the number of female offspring produced annually (Fig 3B). *Rhynchobatus australiae* and *G. cemiculus* had the highest annual reproductive output and *r*_*max*_, while *G. typus* had lower *r*_*max*_ estimates but the same annual reproductive output as the two species (Fig 3B). *Rhinobatos rhinobatos, P. horkelii* and *Z. exasperata* had similar estimates of annual reproduction, yet *R. rhinobatos* had a higher estimates of *r*_*max*_ than *P. horkelii* and *Z. exasperata* (Fig 3B). *Z*. brevirostris had the lowest annual reproductive output and *r*_*max*_ estimate (Fig 3B). Maximum rate of population growth increased with maximum size of the species (Fig 4A). The largest species (i.e. *R. australiae, G. cemiculus* and *G. typus*) were estimated to have a higher maximum rate of population increase than the smaller species in the order, such as *P. horkelii* and *Z. brevirostris* (Table 4; Fig 4A). The largest species, *R. australiae, G. cemiculus* and *G. typus*, had the highest mean annual reproductive output and the largest mean size at birth in relation to the maximum size, compared to the six smaller species (Fig 4B, C). The smallest species, *Z. exasperata* and *Z. brevirostris* had the smallest annual reproductive output and size at birth, in relation to their maximum size (Fig 4B, C).

**Figure 3.**
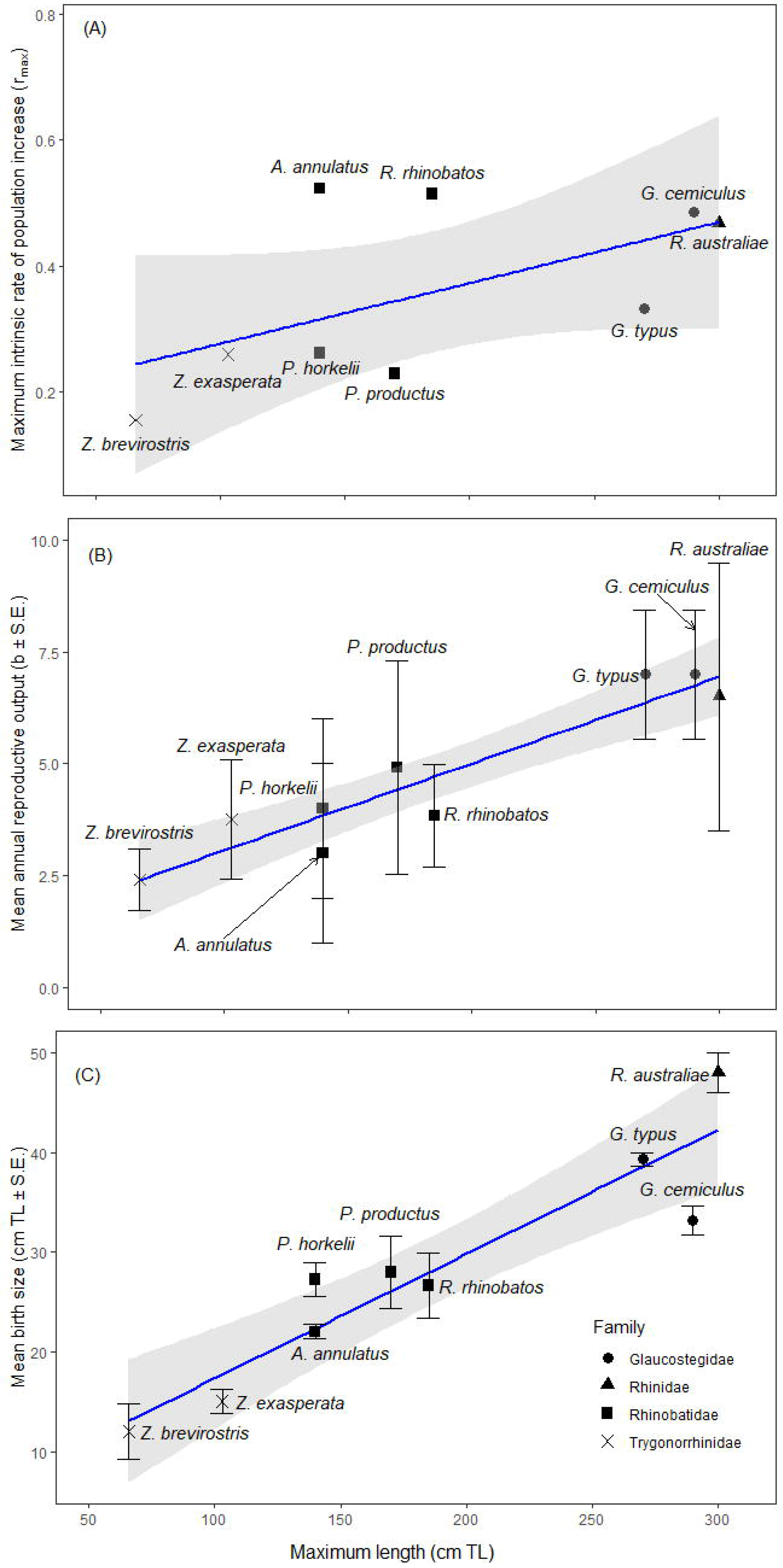
Maximum intrinsic rate of population increase (*r*_*max*_) for the nine species of wedgefishes, guitarfishes, and banjo rays in relation to the (A) mean ± S.E. age at maturity (*a*_*mat*_, years) and (B) mean ± S.E. annual reproductive output (*b*). The shapes represent the four families, black circle represents the Family Glaucostegidae (giant guitarfishes), black triangle signifies the Rhinidae (wedgefishes), black squares represents species from Rhinobatidae (guitarfishes) and black crosses are the species from Trygonorrhinidae (banjo rays).

**Figure 4.**
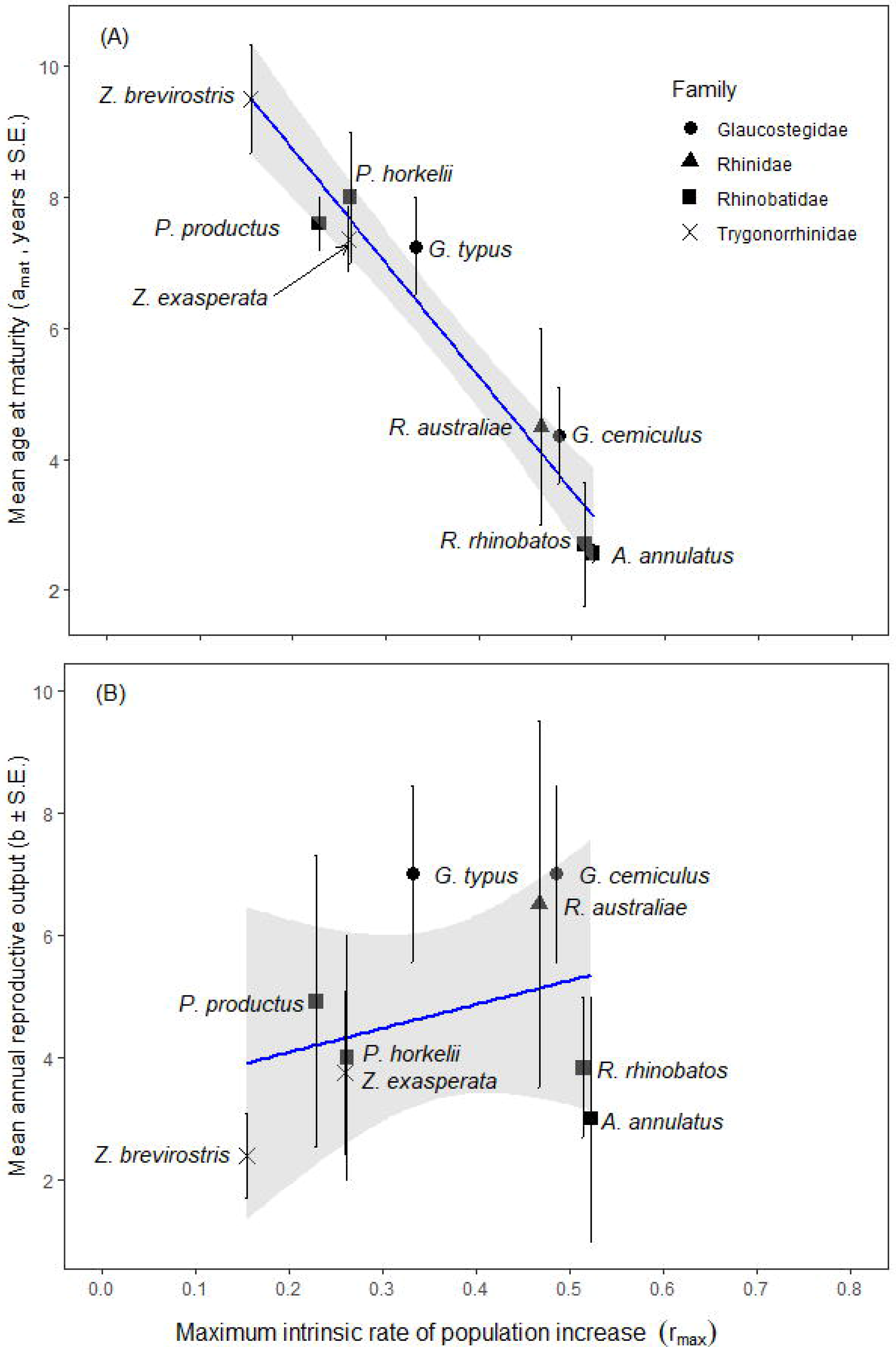
Maximum size (cm TL) for the nine species of wedgefishes, guitarfishes, and banjo rays in relation to the (A) median maximum intrinsic rate of population increase (*r*_*max*_, year^−1^) using the reciprocal of lifespan to estimate natural mortality, (B) mean (± S.E.) annual reproduction rate of females (*b*), and (C) mean (± S.E.) size at birth (cm TL). The shapes represent the four families, black circle represents the Family Glaucostegidae (giant guitarfishes), black triangle signifies the Rhinidae (wedgefishes), black squares represents species from Rhinobatidae (guitarfishes) and black crosses are the species from Trygonorrhinidae (banjo rays).

For the 115 chondrichthyan species, maximum intrinsic rate of population increase ranged from 0.035 year^−1^ for the gulper shark *Centrophorus granulosus* (Family Centrophoridae) to 1.395 year^−1^ for the brown skate *Raja miraletus* (Family Rajidae) (Fig 5). The maximum intrinsic rate of population increase of wedgefishes, guitarfishes, and banjo rays was similar to that of the four sawfish species (Family Pristidae), *Pristis pristis* (0.172 year^−1^), *Pristis pectinate* (0.179 year^−1^), *Pristis clavata* (0.236 year^−1^) and *Pristis zijsron* (0.272 year^−1^). Estimates of *r*_*max*_ for *Acroteriobatus annulatus, R. rhinobatos*, *G. cemiculus* and *R. australiae* were among the highest among chondrichthyans (median = 0.522, 0.539, 0.486 and 0.467 year^−1^, respectively). *Rhynchobatus australiae*, *G. cemiculus* and *G. typus* had relatively high *r*_*max*_ estimates, compared to species with similar maximum sizes (Fig 6A). *Pseudobatos horkelii, P. productus* and *Z. exasperata* had mid-range estimates of *r*_*max*_ compared to species of a similar size (Fig 6A). Compared to other species of the same maximum size, *A. annulatus* and *R. rhinobatos* had relatively high *r*_*max*_, while *Z. brevirostris* had a lower *r*_*max*_ compared to similar sized species (Fig 6A). The majority of the largest chondrichthyan species for which *r*_*max*_ are all listed on CITES and CMS, however they are not the least productive species (Fig 6A).

Chondrichthyans species with the oldest age at maturity tend to have the lowest *r*_*max*_ estimates (Fig 6B). *Acroteriobatus annulatus*, *G*. cemiculus and *R. australiae* mature at the youngest ages and had higher estimates of *r*_*max*_, compared to the other rhinopristiformes and chondrichthyans (Fig 6B). As maximum age increased, *r*_*max*_ estimates decreased for the 115 chondrichthyans species (Fig 6C). *Acroteriobatus annulatus*, *R. rhinobatos, G*. cemiculus and *R. australiae* are among the chondrichthyans species with the lowest maximum age estimates, and hence high *r*_*max*_ (Fig 6C). *Glaucostegus typus, Z. exasperata, P. horkelii* and *P. productus* have mid-range maximum age compared to other species, while *Z. brevirostris* had a lower *r*_*max*_ estimate compared to other species with a similar maximum age (Fig 6C).

Across chondrichthyan species, as the number of female offspring produced annually increases, *r*_*max*_ also increases (Fig 6D). However, *A. annulatus*, *R. rhinobatos, G*. cemiculus and *R. australiae* have a relatively higher *r*_*max*_ estimates compared to species with similar annual reproductive output. *Zapteryx exasperata, P. horkelii* and *P. productus* are estimated to have a mid-range annual reproductive estimate, compared to species with similar *r*_*max*_ (Fig 6D). *Glaucostegus typus* has a relatively high *r*_*max*_ estimate compared to species with similar annual reproductive output, while *Z. brevirostris* has a low *r*_*max*_ estimate compared to species with similar annual reproductive output (Fig 6D). The majority of currently CITES listed species have a low annual reproductive output and *r*_*max*_ (Fig 6D).

There is a positive correlation between growth coefficient and maximum intrinsic rate of population increase (Fig 6E). *Acroteriobatus annulatus*, *R. rhinobatos, G*. cemiculus and *R. australiae* have fast somatic growth and a high *r*_*max*_, in comparison to the other chondrichthyan species (Fig 6E). *Glaucostegus typus, Z. exasperata* and *P. horkelii* have a mid-range *r*_*max*_ estimate compared to species with similar growth rates, while *P. productus* and *Z. brevirostris* have a lower *r*_*max*_ estimate compared to other species with similar growth rates (Fig 6E). The majority of the CITES and CMS listed species have a relatively low growth coefficients and *r*_*max*_ estimates (Fig 6E).

**Figure 5.**
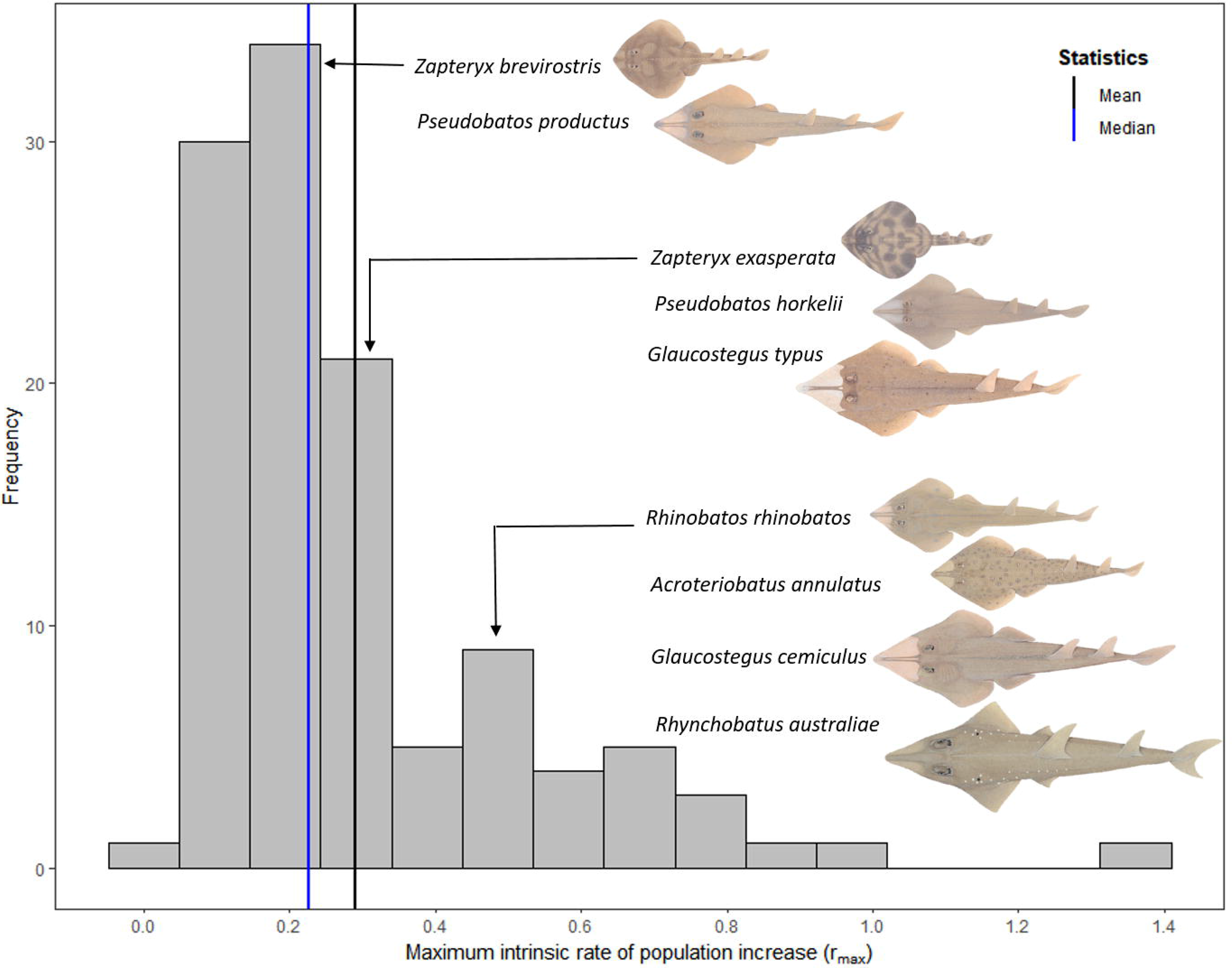
Median maximum intrinsic rate of population increase (*r*_*max*_) for 115 chondrichthyans, including the nine wedgefishes, guitarfishes, and banjo ray species, which are displayed on the figure and grouped by their *r*_*max*_ values. Blank line denote the mean (*r*_*max*_ = 0.30) and blue line represents the median (*r*_*max*_ = 0.23). Species illustrations are from [66]

**Figure 6.**
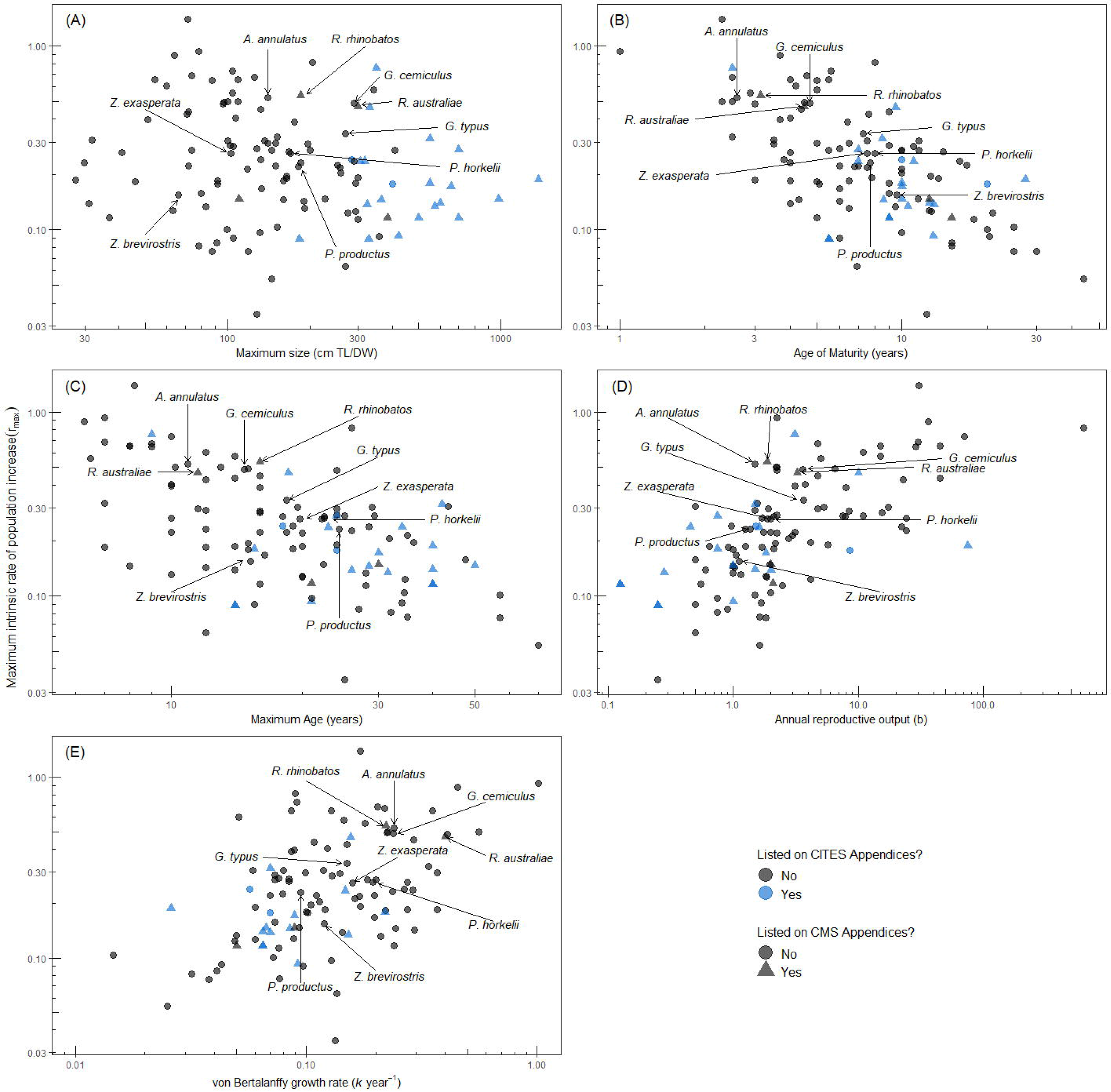
Maximum intrinsic rate of population increase (*r*_*max*_) estimates for 115 chondrichthyans, including the nine wedgefishes, guitarfishes, and banjo ray species compared with (A) maximum size (cm TL/DW/FL), (B), age at maturity (*α*_*mat*_ years), (C) maximum age (*α*_*max*_, years), (D) annual reproductive output *b*, (E) the von Bertanlaffy growth coefficient (*k*, year^−1^). All axes are on a logarithmic scale. The nine wedgefishes, guitarfishes, and banjo ray species are labelled on the graph, and the median *r*_*max*_ value is reported, using the reciprocal of the lifespan method to estimate natural mortality. Species that are listed on CITES appendix I or II represented in blue, non-CITES species in grey, and species listed on CMS appendix I or II represented as triangle, with non-CMS species are closed circle.

## Discussion

Our analysis show that the larger species such as the wedgefishes and giant guitarfishes appear to have a greater intrinsic ability to recover from population depletion than many other large chondrichthyans including those currently list on multilateral environmental agreements like CITES and CMS, such as manta and devil rays [28, 29]. Maximum rate of intrinsic population growth increased with increasing body size for nine species of rhinopristiformes. The largest species, *R. australiae*, *G. cemiculus* and *G. typus* had among the highest estimates of population increase for these rays, while the smaller banjo rays (family Trygonorrhinidae) such as the *Z. brevirostris* and *Z. exasperata* had the lowest estimates. The two species of guitarfish (family Rhinobatidae), *A. annulatus* and *R. rhinobatos*, did not fall within the positive relationship between maximum sizes and *r*_*max*_ seen in this group. *Acroteriobatus annulatus* and *R. rhinobatos* only attaining 140 cm TL and 100 cm TL, respectively [66], and had the highest estimates of *r*_*max*_. The high *r*_*max*_ estimates for *A*. *annulatus* and *R. rhinobatos* could be explained by the young age at maturity (1.9 – 4.5 years), fast somatic growth (0.134 – 0.289 *k* years^−1^), and high annual reproductive output (2.5–12 females pups per year). The high maximum intrinsic rate of population growth for larger species, such as wedgefishes and giant guitarfishes, which both reach maximum sizes close to 300cm TL [66], is the result of the higher annual reproductive outputs, combined with early at age maturity, compared to other species of a similar maximum size. While the banjo rays, *Z. brevirostris* and *Z. exasperata* which reach a maximum size of 66 cm TL and 97cm TL, respectively, had slower growth, later maturity compared to other similar sized chondrichthyans species. However, this trend is highly unusual among elasmobranchs [101]. These findings contrasts other multi-species comparative studies, such as Dulvy et al. (28), where the maximum intrinsic rate tends to decrease with increasing maximum size, however this study had more sharks than large tropical rays. The productivity of wedgefishes, guitarfishes, and banjo rays was more similar to four sawfish species: largetooth sawfish *Pristis pristis* (0.172 year^−1^), smalltooth sawfish *Pristis pectinata* (0.179 year^−1^), dwarf sawfish *Pristis clavata* (0.236 year^−1^) and green sawfish *Pristis zijsron* (0.272 year^−1^). Despite their large size (ranging from 318 – 700 cm TL), some species of sawfish have been estimated to have a relatively high productivity for elasmobranchs [59]. Typically large bodied marine animals are associated with factors of vulnerability such as lower intrinsic rate of population growth, late maturity, and dependence on vulnerable habitat, while smaller bodied species are linked to factors providing resilience including faster population growth and early maturity [14, 27, 76]. While body size has been used to predict extinction risk in elasmobranchs, with the larger species predicted to be most at risk of extinction [16], some studies have found little [9, 15] to no correlation [20] between body size and rate of population increase. The relationship between body size and rate of population growth has been hypothesised to be the result of correlations between body size and other more influential life history traits such as age at maturity and litter size [102, 103], which may be the case for the wedgefishes, guitarfishes, and banjo rays.

Many elasmobranch species have a low reproductive rate [20]. Three species had annual reproductive outputs less than 5 pups per year, *P. productus*, *Z. brevirostris* and *A. annulatus*. Species with reproductive output lower than five pups per year have been demonstrated to have the highest demographic uncertainty when estimating *r*_*max*_, and are likely to have low *r*_*max*_ estimates [14], which was observed for *P. productus* and *Z. brevirostris*. However, while *A. annulatus* had very low annual reproductive output (*b* = 1-5), this species also had one of the highest estimates of *r*_*max*_. The high estimate of *r*_*max*_ may be attributed instead to the very young age at maturity (*α*_*mat*_ = 2-3 years), as well as the fast growth (*k* = 0.240 year^−1^) and long lifespan (*α*_*max*_ = 15 years). The annual reproductive output was higher in the larger species, such as *R. australiae and G. cemiculus*, which have the largest annual reproductive output parameters (*b* = 3.5 − 9.5 and *b* = 2.5 − 12, respectively). This reproductive output is likely to be driving the high *r*_*max*_ estimates for these species. While the similar sized *G. typus* had the same annual reproductive output as *G. cemiculus*, it has lower estimate of *r*_*max*_, likely the result of an older age at maturity (*α*_*mat*_ = 6.5 − 8 years) than *R. australiae* (*α*_*mat*_ = 3 − 6 years) and *G. cemiculus* (*α*_*mat*_ = 2.9 − 6.5 years). As *r*_*max*_ estimates are sensitive to increasing variation in age at maturity [14], the early maturity of many wedgefishes, guitarfishes, and banjo rays, particularly compared to species of similar sizes, may help to explain the relatively high *r*_*max*_ estimates for this group. The larger body size of wedgefish and giant guitarfish species (max. size estimated between 270-300 cm TL) allows these species to produce numerous and large offspring in relation to their maximum size. Whereas the guitarfishes and banjo rays have smaller birth size and smaller litters relative to their maximum size. Larger offspring will likely have a greater survival probability than the smaller offspring of species with a similar *r*_*max*_ [14], and for long lived species, juvenile survival is a key contributor to the population growth rate [9]. While the model used in this study incorporates juvenile survival, it also assumes that juvenile mortality is equal to adult mortality [64]. As juveniles tend to have higher mortality rates than adults [104], which then can vary with local differences in habitat [105], this assumption of equal mortality is likely to result in conservative estimates of *M* [64]. The differential juvenile mortality among species was not accounted for in this model but should be the focus of further study [14].

Natural mortality, referring to the death of individuals in the population from natural causes such as predation, disease and old age [94], is one of the most important parameters in fisheries and conservation modelling, yet it is one of the hardest to estimate [10, 106, 107]. While in some models uncertainty in the natural mortality parameter has had little influence on *r*_*max*_ [14], the different estimators can have substantial effects on *r*_*max*_ estimates [107]. There is considerable debate as to which empirical model should be used to estimate adult natural mortality, as there are numerous and diverse approaches using life history information to estimate this parameter [106, 108]. Jensen’s First Estimator [98] and the modified Hewitt and Hoeing method [99] systematically resulted in negative, and thus unrealistic estimates, of *r*_*max*_ for five out of the nine species of shark-like ray species, which was also demonstrated in [64], and were considered to be less appropriate for this group. The implausible estimates are likely the consequence of overestimating natural mortality (e.g. > 0.1 year^−1^) for these species, particularly when the annual reproductive output is low (e.g. b < 5) and age at maturity is high [14, 64]. The other two natural mortality estimators, the Frisk’s estimator and Reciprocal of lifespan, provided similar estimates of maximum rate of population increase for *G. cemiculus, G. typus, R. rhinobatus, P. horkelii, Z. exasperata* and *Z. brevirostris*. The plausible *r*_*max*_ estimates produced by these two natural mortalities suggest these natural mortality estimators may be the more appropriate methods for elasmobranchs. Frisk’s estimator and Reciprocal of life span are more suited for elasmobranchs, given they have a relatively high juvenile survival [9, 64], while the Jensen’s First Estimator and the modified Hewitt and Hoeing method were explicitly designed for adult mortality. Most teleost species have different life history strategies to chondrichthyan species (i.e. higher fecundity, younger age at maturity, faster growth, and shorter lifespans) [109, 110]. Thus, it is likely that Jensen’s First Estimator and the modified Hewitt and Hoeing are less appropriate methods of estimating natural mortality for chondrichthyans. Unfortunately estimating natural mortality with useful accuracy is remarkably difficult [108]. Identifying, or improving the best indirect estimator would require data-intensive methods, such as catch data to analyse catch curves, mark re-capture experiments, virtual population analysis (VPA), or fully integrated stock assessments [108]. These methods all require extensive prior knowledge of the species biology which is lacking for many chondrichthyan species. Presenting the results from multiple natural mortality estimators provides a better understanding of the uncertainty associated with the maximum intrinsic rate of population increase.

The greatest obstacle to accurately estimate *r*_*max*_ and natural mortality is the accuracy of the biological information used [91]. The use of inaccurate surrogate information can reduce the accuracy of the demographic models [91, 111, 112]. Of the 56 species across the four families of wedgefishes, guitarfishes, and banjo rays, only nine species had sufficient information to estimate their maximum intrinsic rate of population increase, and with relatively high levels of uncertainty associated with the life history parameters. For example, there were only age and growth two studies for wedgefishes and giant guitarfishes, one from the eastern coast of Australia for *R. australiae* and *G. typus* [65], and one from Central Mediterranean Sea for *G. cemiculus* [70]. Neither study estimated age at maturity, nor aged individuals at the maximum sizes. Given that the age at maturity is a pivotal parameter when estimating *r*_*max*_, yet highly uncertain for all wedgefishes, guitarfishes, and banjo rays examined, these estimates must be taken with caution. Furthermore, numerous reviews have reported sampling biases and failures in ageing protocols, including lack of validation [113, 114] that often result in overestimation or underestimate of age and growth parameters [115]. As there has been no validation studies in the ages of wedgefishes, guitarfishes, and banjo rays, the maximum ages for these species are likely to be underestimated, while the age at maturity estimates could also be inaccurate. This can lead to inaccurate estimates of natural mortality and *r*_*max*_ [91, 116]. The information on the reproductive biology for rhinopristiforms is generally limited, but is more available for species in the Rhinobatidae and Trygonorrhinidae families. For example, there is evidence that species such as *P. productus, P. horkelii*, and Z*. exasperata* experience embryonic diapause or delayed development [87, 117], potentially as a result of unfavourable environmental conditions [118] or sex segregation [119]. In addition, Simpfendorfer (120) hypothesised that diapause allowed other elasmobranch species (*Rhizoprionodon taylori*) to have larger litter sizes than other similar sized species in the same family (Carcharhinidae). As possibility of diapause was not able to be taken into account during this study, the breeding interval and annual reproductive output may be inaccurate for *Z. exasperata*, thus resulting in an inappropriate maximum intrinsic rate of population growth. Directing research efforts to obtain data from more species, as well as improving the accuracy of life history parameters for data poor species, such as age at maturity and annual reproductive output, would be the most pragmatic option to improve the accuracy of *r*_*max*_ for wedgefishes, guitarfishes, and banjo rays.

Measuring the population productivity of a species allows for a greater understanding of the species’ ability to recover from declines, and has direct practical importance in conservation and fisheries [91, 121]. Compared to other chondrichthyans, wedgefishes, giant guitarfishes and guitarfishes appear to have a high maximum intrinsic rate of population increase, therefore possessing biological traits that can potentially allow them to recover from population declines more rapidly than other threatened species. While other families, such as banjo rays, are less productive and may have a slower ability to rebound from population depletions. However, the current and unregulated fishing pressure that most wedgefishes, guitarfishes, and banjo ray species currently experience is likely unsustainable, with reported severe declines and localised extinctions of wedgefishes and guitarfishes throughout their range [34, 48]. Therefore, it appears that these groups are facing a widespread conservation crisis [34]. Yet globally, wedgefishes, guitarfishes, and banjo rays are currently not managed through international trade or fishing restrictions, and there is limited market and trade information available [122]. Given the lack of national level management for these species, and the role of international trade, especially in fins, the use of trade controls through CITES listings may be a useful way encouraging better management of their wedgefishes, guitarfishes, and banjo ray species. Currently, the families Rhinidae and Glaucostegidae are being proposed for CITES Appendix II listing, which are for species that not necessarily currently threatened with extinction but may become so unless trade is controlled [123]. However, trade restrictions alone will not reduce fishing mortality since these species are rarely taken in directed fisheries (although there are exceptions e.g. in Indonesia [43]). Regional and national fisheries management strategies will be required to address the overfishing of stocks, which will require reductions in fishing effort. Lastly, increasing the accuracy of information and research on the biology and life history of these species will be essential for their management and conservation.

## Conclusion

Using current life history data, incorporating uncertainty in parameters, and taking into account juvenile mortality, this study provides the first analysis into the population productivity for nine species from four families of rhinopristiforms. The larger wedgefishes and giant guitarfishes were found to be relative productive families, while the smaller guitarfishes and banjo rays were less productive, compared to other chondrichthyans. While the maximum intrinsic rate of population growth varied with the different natural mortality estimator, *r*_*max*_ appears to increase with increasing maximum size for the four families, which is counter to most studies of shark populations. There was considerable uncertainty in the age at maturity and annual reproductive output for all species. There is a need for accurate life history information for these data poor species, as well directing efforts to obtain data for more species as there was only nine of out 56 species with sufficient life history information. We recommend presenting the results from multiple natural mortality estimators to provide a greater understanding of the uncertainty for the maximum intrinsic rate of population increase. It appears that wedgefishes and giant guitarfishes could recover from population depletion faster than guitarfishes and banjo rays, if fishing mortality is kept low. Extensive regional, national and international management strategies will be required to address the overfishing of stock and reduction of fishing mortality, and the use of international trade regulations such as CITES listing, may help to achieve positive conservation outcomes. The results of this study emphasise the urgency for effective management for the conservation of these rays.

## Acknowledgements

The authors would like to thank Charlotte Heacock for assisting in the data collection. This project was funded by the Shark Conservation Fund (BMD), a philanthropic collaborative pooling expertise and resources to meet the threats facing the world’s sharks and rays. The Shark Conservation Fund is
a project of Rockefeller Philanthropy Advisors. The corresponding author (BMD) is supported through an Australian Government Research Training Program Scholarship (RTPS). The scientific results and conclusions, as well as any views or opinions expressed herein, are those of the author(s) and do not necessarily reflect those of NOAA or the Department of Commerce. The funders had no role in study design, data collection and analysis, decision to publish, or preparation of the manuscript.

## Author contribution

**Conceptualisation:** Brooke M. D’Alberto, John K. Carlson, Colin A. Simpfendorfer,

**Data curation:** Brooke M. D’Alberto

**Data analysis:** Brooke M. D’Alberto

**Funding acquisition:** Brooke M. D’Alberto

**Investigation:** Brooke M. D’Alberto

**Methodology:** Sebastian A. Pardo, Brooke M. D’Alberto

**Project administration:** Brooke M. D’Alberto

**Visualisation:** Brooke M. D’Alberto, Sebastian A. Pardo, Colin A. Simpfendorfer

**Writing – original draft:** Brooke M. D’Alberto

**Writing – reviewing & editing:** John K. Carlson, Sebastian Pardo, Colin A. Simpfendorfer

## Supporting Information

**S1 Table.** Maximum intrinsic rate of population increase (*r*_*max*_) estimates, life history values and sources used to estimate *r*_*max*_ for 115 chondrichthyans, including the nine wedgefishes, guitarfishes, and banjo ray species, using the reciprocal of the lifespan method to estimate natural mortality. The values included are the maximum size (Max. size in centimetres total length/disk width, cm TL/DW), von Bertalanffy growth coefficient (*k*, year^−1^), age at maturity (*α*_*mat*_, years), reported maximum age (*α*_*max*_, years), litter size (*l*), breeding interval (*i*, years), annual reproductive output of females (*b*). Included is whether the species are listed on the appendixes of Convention of International Trade of Endangered Species (CITES, yes or no) and/or Convention on the Conservation of Migratory Species of Wild Animals (CMS, yes or no). The ‘na’ indicates parameter was not available from literature.

**S2 Figure.** Predicted values of maximum intrinsic rate of population increase (*r*_*max*_) for nine wedgefishes, guitarfishes, and banjo ray species when including uncertainty in the Jensen’s natural mortality estimator (*M*, top/grey boxplot), annual reproductive output (*b*, middle/blue boxplot), and age at maturity (*α*_*mat*_, bottom/orange boxplot). Species are (A) *Rhynchobatus australiae*, (B), *Glaucostegus cemiculus*, (C) *Glaucostegus typus*, (D) *Acroteriobatus annulatus*, (E) *Pseudobatos horkelii*, (F) *Pseudobatos productus*, (G) *Rhinobatos rhinobatos*, (H) *Zapteryx brevirostris*, and (I) *Zapteryx exasperata*. Boxes indicate median, 25 and 75% quantiles, whereas the lines encompass 95% of the values (2.5 and 97.5% quantiles).

**S3 Figure.** Predicted values of maximum intrinsic rate of population increase (*r*_*max*_) for nine wedgefishes, guitarfishes, and banjo ray species when including uncertainty in the modified Howitt & Hewitt’s natural mortality estimator (*M*, top/grey boxplot), annual reproductive output (*b*, middle/blue boxplot), and age at maturity (*α*_*mat*_, bottom/orange boxplot). Species are (A) *Rhynchobatus australiae*, (B), *Glaucostegus cemiculus*, (C) *Glaucostegus typus*, (D) *Acroteriobatus annulatus*, (E) *Pseudobatos horkelii*, (F) *Pseudobatos productus*, (G) *Rhinobatos rhinobatos*, (H) *Zapteryx brevirostris*, and (I) *Zapteryx exasperata*. Boxes indicate median, 25 and 75% quantiles, whereas the lines encompass 95% of the values (2.5 and 97.5% quantiles).

**S4 Figure.** Predicted values of maximum intrinsic rate of population increase (*r*_*max*_) for nine wedgefishes, guitarfishes, and banjo ray species when including uncertainty in the Frisk’s natural mortality estimator (*M*, top/grey boxplot), annual reproductive output (*b*, middle/blue boxplot), and age at maturity (*α*_*mat*_, bottom/orange boxplot). Species are (A) *Rhynchobatus australiae*, (B), *Glaucostegus cemiculus*, (C) *Glaucostegus typus*, (D) *Acroteriobatus annulatus*, (E) *Pseudobatos horkelii*, (F) *Pseudobatos productus*, (G) *Rhinobatos rhinobatos*, (H) *Zapteryx brevirostris*, and (I) *Zapteryx exasperata*. Boxes indicate median, 25 and 75% quantiles, whereas the lines encompass 95% of the values (2.5 and 97.5% quantiles).

