## Supplementary material for "Population productivity of wedgefishes, guitarfishes, and banjo rays: inferring the potential for recovery": S1 Table

| **Order** | **Family** | **Species** | **CITES** | **CMS** | **Max. size (cm TL/DW)** | ***k* (year^-1^)** | ***α_mat_* (years)** | ***α_max_* (years)** | **average lifespan** | ***l*** | ***i* (years)** | ***b*** | ***r_max_*** | **Source** |
| --- | --- | --- | --- | --- | --- | --- | --- | --- | --- | --- | --- | --- | --- | --- |
| Carchariniformes | Carcharhinidae | *Carcharhinus acronotus* | No | No | 132.4 | 0.266 | 3.8 | 19.0 | 11.4 | 3.9 | 2 | 1.03 | 0.241 | [1] |
| Carchariniformes | Carcharhinidae | *Carcharhinus amblyrhynchos* | No | No | 190.0 | 0.294 | 6.0 | 12.0 | 9.0 | 4.1 | 2 | 1.00 | 0.142 | [1] |
| Carchariniformes | Carcharhinidae | *Carcharhinus brachyurus* | No | No | 275.0 | 0.049 | 20.9 | 34.5 | 27.7 | 16.6 | 2 | 1.98 | 0.124 | [1] |
| Carchariniformes | Carcharhinidae | *Carcharhinus brevipinna* | No | No | 294.1 | 0.100 | 9.0 | 19.0 | 14.0 | 8.5 | 2 | 1.05 | 0.180 | [1] |
| Carchariniformes | Carcharhinidae | *Carcharhinus cautus* | No | No | 133.0 | 0.198 | 6.0 | 16.4 | 11.2 | 4.2 | 2 | 8.63 | 0.167 | [1] |
| Carchariniformes | Carcharhinidae | *Carcharhinus falciformis* | Yes | Yes | 305.0 | 0.148 | 9.0 | 17.9 | 13.5 | 6.5 | 2 | 1.67 | 0.155 | [1] |
| Carchariniformes | Carcharhinidae | *Carcharhinus galapagensis* | No | No | 300.0 | 0.172 | 7.8 | 15.0 | 11.4 | 8.7 | 2 | 2.13 | 0.193 | [1] |
| Carchariniformes | Carcharhinidae | *Carcharhinus isodon* | No | No | 159.6 | 0.244 | 4.3 | 8.0 | 6.2 | 4.0 | 2 | 1.25 | 0.146 | [1] |
| Carchariniformes | Carcharhinidae | *Carcharhinus leucas* | No | No | 300.2 | 0.076 | 18.0 | 28.0 | 23.0 | 9.9 | 2 | 5.75 | 0.114 | [1] |
| Carchariniformes | Carcharhinidae | *Carcharhinus limbatus* | No | No | 191.0 | 0.210 | 6.5 | 10.0 | 8.3 | 4.6 | 2 | 2.18 | 0.131 | [1] |
| Carchariniformes | Carcharhinidae | *Carcharhinus longimanus* | Yes | No | 285.0 | 0.101 | 5.5 | 14.0 | 9.8 | 6.1 | 2 | 1.85 | 0.212 | [1] |
| Carchariniformes | Carcharhinidae | *Carcharhinus obscurus* | No | No | 357.2 | 0.043 | 20.5 | 34.0 | 27.3 | 10.0 | 3 | 4.15 | 0.092 | [1] |
| Carchariniformes | Carcharhinidae | *Carcharhinus plumbeus* | No | No | 226.5 | 0.093 | 11.4 | 22.4 | 16.9 | 7.9 | 2 | 2.78 | 0.148 | [1] |
| Carchariniformes | Carcharhinidae | *Carcharhinus porosus* | No | No | 128.0 | 0.076 | 6.0 | 24.0 | 15.0 | 4.5 | 1 | 1.63 | 0.275 | [1] |
| Carchariniformes | Carcharhinidae | *Carcharhinus signatus* | No | No | 260.0 | 0.114 | 10.0 | 31.7 | 20.9 | 11.1 | 2 | 0.98 | 0.204 | [1] |
| Carchariniformes | Carcharhinidae | *Carcharhinus sorrah* | No | No | 151.8 | 0.340 | 2.5 | 7.0 | 4.8 | 3.1 | 1 | 1.53 | 0.320 | [1] |
| Carchariniformes | Carcharhinidae | *Carcharhinus tilstoni* | No | No | 196.0 | 0.140 | 3.5 | 12.0 | 7.8 | 3.0 | 1 | 7.88 | 0.292 | [1] |
| Carchariniformes | Carcharhinidae | *Galeocerdo cuvier* | No | No | 410.0 | 0.184 | 10.0 | 22.5 | 16.3 | 31.5 | 2 | 7.50 | 0.271 | [1] |
| Carchariniformes | Carcharhinidae | *Isogomphodon oxyrhynchus* | No | No | 160.0 | 0.121 | 6.5 | 20.0 | 13.3 | 5.0 | 2 | 2.25 | 0.185 | [1] |
| Carchariniformes | Carcharhinidae | *Negaprion brevirostris* | No | No | 293.6 | 0.060 | 12.7 | 20.0 | 16.4 | 7.4 | 2 | 1.50 | 0.126 | [1] |
| Carchariniformes | Carcharhinidae | *Prionace glauca* | No | No | 341.8 | 0.146 | 5.0 | 14.0 | 9.5 | 30.0 | 1 | 4.75 | 0.577 | [1] |
| Carchariniformes | Carcharhinidae | *Rhizoprionodon taylori* | No | No | 78.4 | 1.013 | 1.0 | 7.0 | 4.0 | 4.5 | 1 | 5.35 | 0.929 | [1] |
| Carchariniformes | Carcharhinidae | *Rhizoprionodon terraenovae* | No | No | 108.9 | 0.560 | 2.3 | 10.2 | 6.3 | 4.4 | 1 | 1.50 | 0.499 | [1] |
| Carchariniformes | Scyliorhinidae | *Scyliorhinus canicula* | No | No | 71.0 | 0.150 | 7.6 | 12.0 | 9.8 | 45.5 | 1 | 1.55 | 0.426 | [1] |
| Carchariniformes | Sphyrnidae | *Sphyrna lewini* | Yes | Yes | 331.0 | 0.156 | 5.8 | 18.6 | 12.2 | 20.1 | 1 | 2.48 | 0.465 | [1] |
| Carchariniformes | Sphyrnidae | *Sphyrna mokarran* | Yes | Yes | 550.0 | 0.070 | 8.5 | 42.0 | 25.2 | 6.0 | 2 | 4.15 | 0.314 | [2, 3] |
| Carchariniformes | Sphyrnidae | *Sphyrna tiburo* | No | No | 104.0 | 0.180 | 2.9 | 6.5 | 4.7 | 10.0 | 1 | 8.00 | 0.557 | [1] |
| Carchariniformes | Sphyrnidae | *Sphyrna zygaena* | Yes | No | 400.0 | 0.070 | 20.0 | 24.0 | 22.0 | 17.3 | 1 | 1.15 | 0.177 | [4] |
| Carchariniformes | Triakidae | *Furgaleus macki* | No | No | 150.0 | 0.369 | 6.5 | 11.5 | 9.0 | 19.0 | 2 | 3.75 | 0.297 | [1] |
| Carchariniformes | Triakidae | *Galeorhinus galeus* | No | No | 164.5 | 0.105 | 12.6 | 36.2 | 24.4 | 24.9 | 3 | 22.75 | 0.196 | [1] |
| Carchariniformes | Triakidae | *Mustelus antarcticus* | No | No | 175.0 | 0.086 | 6.4 | 16.0 | 11.2 | 16.0 | 1 | 4.75 | 0.387 | [1] |
| Carchariniformes | Triakidae | *Mustelus californicus* | No | No | 125.0 | 0.218 | 2.5 | 9.0 | 5.8 | 9.5 | 1 | 10.05 | 0.673 | [1] |
| Carchariniformes | Triakidae | *Mustelus canis* | No | No | 132.0 | 0.292 | 4.4 | 16.0 | 10.2 | 9.5 | 1 | 2.20 | 0.452 | [1] |
| Carchariniformes | Triakidae | *Mustelus henlei* | No | No | 100.0 | 0.225 | 2.5 | 13.0 | 7.8 | 4.5 | 1 | 2.25 | 0.500 | [1] |
| Carchariniformes | Triakidae | *Mustelus lenticulatus* | No | No | 137.0 | 0.119 | 7.5 | 19.5 | 13.5 | 10.7 | 1 | 5.00 | 0.304 | [1] |
| Carchariniformes | Triakidae | *Mustelus manazo* | No | No | 107.0 | 0.124 | 4.0 | 10.0 | 7.0 | 7.5 | 1 | 15.00 | 0.404 | [1] |
| Carchariniformes | Triakidae | *Mustelus mustelus* | No | No | 164.0 | 0.060 | 13.5 | 24.0 | 18.8 | 11.5 | 1 | 4.75 | 0.190 | [1] |
| Carchariniformes | Triakidae | *Triakis semifasciata* | No | No | 145.0 | 0.073 | 10.0 | 25.0 | 17.5 | 15.0 | 1 | 2.25 | 0.272 | [1] |
| Chimaeriformes | Callorhinchidae | *Callorhinchus capensis* | No | No | 60.0 | 0.051 | 4.2 | 12.0 | 8.1 | 22.0 | 1 | 11.00 | 0.604 | [1] |
| Chimaeriformes | Callorhinchidae | *Callorhinchus milii* | No | No | 97.0 | 0.224 | 4.5 | 15.0 | 9.8 | 13.0 | 1 | 6.50 | 0.492 | [1] |
| Chimaeriformes | Chimaeridae | *Chimaera monstrosa* | No | No | 74.0 | 0.084 | 11.5 | 29.4 | 20.5 | 22.0 | 1 | 11.00 | 0.272 | [1] |
| Hexanchiformes | Hexanchidae | *Notorynchus cepedianus* | No | No | 291.0 | 0.291 | 16.0 | 28.5 | 22.3 | 88.5 | 2 | 22.13 | 0.236 | [1] |
| Lamniformes | Alopiidae | *Alopias pelagicus* | Yes | Yes | 365.2 | 0.085 | 8.6 | 28.5 | 18.6 | 2.0 | 1 | 0.50 | 0.145 | [1] |
| Lamniformes | Alopiidae | *Alopias superciliosus* | Yes | Yes | 422.0 | 0.092 | 12.9 | 21.0 | 17.0 | 2.0 | 1 | 1.00 | 0.093 | [1] |
| Lamniformes | Alopiidae | *Alopias vulpinus* | Yes | Yes | 573.0 | 0.153 | 10.5 | 31.5 | 21.0 | 1.1 | 2 | 2.08 | 0.134 | [5-7] |
| Lamniformes | Cetorhinidae | *Cetorhinus maximus* | Yes | Yes | 980.0 | 0.067 | 10.0 | 50.0 | 30.0 | 6.0 | 3 | 1.50 | 0.147 | [1] |
| Lamniformes | Lamnidae | *Carcharodon carcharias* | Yes | Yes | 600.0 | 0.065 | 12.5 | 40.0 | 26.3 | 6.0 | 2 | 0.28 | 0.140 | [1] |
| Lamniformes | Lamnidae | *Isurus oxyrinchus* | No | Yes | 385.0 | 0.050 | 15.0 | 21.0 | 18.0 | 12.5 | 3 | 1.00 | 0.116 | [1] |
| Lamniformes | Lamnidae | *Lamna ditropis* | No | No | 257.3 | 0.170 | 7.5 | 20.0 | 13.8 | 4.5 | 1 | 1.00 | 0.219 | [1] |
| Lamniformes | Lamnidae | *Lamna nasus* | Yes | Yes | 324.3 | 0.070 | 13.0 | 26.0 | 19.5 | 4.0 | 1 | 2.00 | 0.137 | [1] |
| Lamniformes | Odontaspididae | *Carcharias taurus* | No | No | 269.5 | 0.136 | 6.9 | 12.0 | 9.5 | 2.0 | 2 | 2.25 | 0.063 | [1] |
| Myliobatiformes | Dasyatidae | *Hypanus americanus* | No | No | 200.0 | 0.200 | 5.5 | 18.0 | 11.8 | 4.2 | 1 | 0.13 | 0.270 | [1] |
| Myliobatiformes | Dasyatidae | *Dasyatis chrysonota* | No | No | 71.9 | 0.070 | 7.0 | 10.0 | 8.5 | 6.2 | 1 | 0.55 | 0.222 | [1] |
| Myliobatiformes | Dasyatidae | *Hypanus dipterurus* | No | No | 83.0 | 0.050 | 9.5 | 28.0 | 18.8 | 2.0 | 1 | 1.00 | 0.133 | [1] |
| Myliobatiformes | Dasyatidae | *Dasyatis pastinaca* | No | No | 51.0 | 0.089 | 3.7 | 10.0 | 6.9 | 6.2 | 1 | 0.60 | 0.395 | [1] |
| Myliobatiformes | Dasyatidae | *Maculabatis astra* | No | No | 80.0 | 0.073 | 9.0 | 47.5 | 28.2 | 1.0 | 1 | 0.65 | 0.158 | [8, 9] |
| Myliobatiformes | Dasyatidae | *Neotrygon picta* | No | No | 32.0 | 0.080 | 3.5 | 43.3 | 23.4 | 1.0 | 1 | 1.75 | 0.306 | [10, 11] |
| Myliobatiformes | Dasyatidae | *Pteroplatytrygon violacea* | No | No | 96.0 | 0.410 | 3.0 | 24.0 | 13.5 | 4.5 | 1 | 0.50 | 0.481 | [1] |
| Myliobatiformes | Mobulidae | *Mobula alfredi* | Yes | Yes | 500.0 | 0.065 | 9.0 | 40.0 | 24.5 | 0.4 | 1.5 | 0.25 | 0.116 | [12] |
| Myliobatiformes | Mobulidae | *Mobula birostris* | Yes | Yes | 700.0 | 0.065 | 9.0 | 40.0 | 24.5 | 0.4 | 1.5 | 0.25 | 0.116 | [12] |
| Myliobatiformes | Mobulidae | *Mobula tarapacana* | Yes | Yes | 328.0 | na | 5.5 | 14.0 | 9.8 | 0.5 | 1 | 0.50 | 0.089 | [12, 13] |
| Myliobatiformes | Mobulidae | *Mobula thurstoni* | Yes | Yes | 183.0 | na | 5.5 | 14.0 | 9.8 | 0.5 | 1 | 0.13 | 0.089 | [14] |
| Myliobatiformes | Myliobatidae | *Aetobatus flagellum* | No | No | 150.0 | 0.111 | 6.0 | 16.0 | 11.0 | 3.5 | 1 | 3.10 | 0.222 | [1] |
| Myliobatiformes | Myliobatidae | *Myliobatis californicus* | No | No | 140.0 | 0.099 | 5.0 | 24.0 | 14.5 | 3.8 | 1 | 1.25 | 0.296 | [1] |
| Myliobatiformes | Rhinopteridae | *Rhinoptera bonasus* | No | No | 104.8 | 0.097 | 6.0 | 15.5 | 10.8 | 1.0 | 1 | 2.10 | 0.090 | [1] |
| Myliobatiformes | Urolophidae | *Trygonoptera mucosa* | No | No | 36.9 | 0.241 | 5.0 | 16.0 | 10.5 | 1.1 | 1 | 1.90 | 0.116 | [1] |
| Myliobatiformes | Urolophidae | *Trygonoptera personata* | No | No | 31.1 | 0.143 | 4.0 | 14.0 | 9.0 | 1.2 | 1 | 0.50 | 0.139 | [1] |
| Myliobatiformes | Urolophidae | *Urolophus lobatus* | No | No | 27.7 | 0.369 | 3.0 | 14.0 | 8.5 | 1.3 | 1 | 3.10 | 0.186 | [1] |
| Myliobatiformes | Urolophidae | *Urolophus paucimaculatus* | No | No | 29.8 | 0.237 | 4.0 | 12.0 | 8.0 | 2.5 | 1 | 2.25 | 0.232 | [1] |
| Orectolobiformes | Rhincodontidae | *Rhincodon typus* | Yes | Yes | 1370.0 | 0.026 | 27.4 | 40.0 | 33.7 | 300.0 | 2 | 75.00 | 0.188 | [1] |
| Rajiformes | Rajidae | *Amblyraja radiata* | No | No | 105.0 | 0.130 | 11.0 | 16.0 | 13.5 | 31.0 | 1 | 24.35 | 0.284 | [1] |
| Rajiformes | Rajidae | *Zearaja chilensis* | No | No | 168.0 | 0.084 | 14.0 | 22.5 | 18.3 | 48.2 | 1 | 24.10 | 0.265 | [1] |
| Rajiformes | Rajidae | *Dipturus trachydermus* | No | No | 253.0 | 0.079 | 17.0 | 26.0 | 21.5 | 48.7 | 1 | 15.50 | 0.226 | [1] |
| Rajiformes | Rajidae | *Leucoraja erinacea* | No | No | 54.0 | 0.352 | 4.0 | 8.0 | 6.0 | 30.0 | 1 | 17.50 | 0.655 | [1] |
| Rajiformes | Rajidae | *Leucoraja naevus* | No | No | 72.0 | 0.108 | 9.0 | 14.0 | 11.5 | 90.0 | 1 | 45.00 | 0.436 | [1] |
| Rajiformes | Rajidae | *Leucoraja ocellata* | No | No | 100.0 | 0.059 | 11.5 | 29.0 | 20.3 | 35.0 | 1 | 28.75 | 0.305 | [1] |
| Rajiformes | Rajidae | *Raja asterias* | No | No | 64.0 | 0.454 | 3.7 | 6.3 | 5.0 | 73.0 | 1 | 45.00 | 0.884 | [1] |
| Rajiformes | Rajidae | *Raja binoculata* | No | No | 203.9 | 0.090 | 8.0 | 26.0 | 17.0 | 1260.0 | 1 | 15.00 | 0.814 | [1] |
| Rajiformes | Rajidae | *Raja brachyura* | No | No | 109.0 | 0.129 | 5.5 | 8.0 | 6.8 | 90.0 | 1 | 30.00 | 0.652 | [1] |
| Rajiformes | Rajidae | *Raja clavata* | No | No | 104.4 | 0.091 | 5.6 | 10.0 | 7.8 | 142.0 | 1 | 71.00 | 0.731 | [1] |
| Rajiformes | Rajidae | *Raja microocellata* | No | No | 87.5 | 0.086 | 5.0 | 9.0 | 7.0 | 57.5 | 1 | 630.00 | 0.649 | [1] |
| Rajiformes | Rajidae | *Raja miraletus* | No | No | 71.7 | 0.172 | 2.3 | 8.2 | 5.3 | 61.0 | 1 | 36.50 | 1.395 | [1] |
| Rajiformes | Rajidae | *Raja montagui* | No | No | 74.0 | 0.204 | 4.6 | 7.0 | 5.8 | 60.0 | 1 | 30.50 | 0.687 | [1] |
| Rhinopristiformes | Glaucostegidae | *Glaucostegus cemiculus* | No | No | 290.0 | 0.237 | 4.7 | 14.7 | 9.7 | 7.0 | 1 | 1.88 | 0.486 | [15-19] |
| Rhinopristiformes | Glaucostegidae | *Glaucostegus typus* | No | No | 270.0 | 0.150 | 7.3 | 18.4 | 12.9 | 7.0 | 1 | 0.75 | 0.333 | [15, 20, 21] |
| Rhinopristiformes | Pristidae | *Anoxypristis cuspidata* | Yes | Yes | 350.0 | na | 2.5 | 9.0 | na | 6.2 | 1 | 0.75 | 0.757 | [22] |
| Rhinopristiformes | Pristidae | *Pristis clavata* | Yes | Yes | 318.0 | na | 7.0 | 34.0 | na | 1.8 | 2 | 1.38 | 0.236 | [22] |
| Rhinopristiformes | Pristidae | *Pristis pectinata* | Yes | Yes | 550.0 | 0.219 | 10.0 | 15.5 | 12.7 | 3.0 | 2 | 1.83 | 0.180 | [22] |
| Rhinopristiformes | Pristidae | *Pristis pristis* | Yes | Yes | 656.0 | 0.089 | 10.0 | 30.0 | 20.0 | 7.3 | 2 | 1.12 | 0.172 | [1] |
| Rhinopristiformes | Pristidae | *Pristis zijsron* | Yes | Yes | 700.0 | na | 7.0 | 24.0 | na | 3.0 | 2 | 0.46 | 0.272 | [22] |
| Rhinopristiformes | Rhinidae | *Rhynchobatus australiae* | No | Yes | 300.0 | 0.400 | 4.5 | 11.5 | 8.0 | 7.0 | 1 | 1.88 | 0.468 | [15, 21] |
| Rhinopristiformes | Rhinobatidae | *Acroteriobatus annulatus* | No | No | 140.0 | 0.240 | 2.6 | 10.9 | 6.8 | 3.0 | 1 | 3.63 | 0.522 | [15, 23] |
| Rhinopristiformes | Rhinobatidae | *Pseudobatos horkelii* | No | No | 170.0 | 0.194 | 8.0 | 22.2 | 15.1 | 4.0 | 1 | 3.25 | 0.261 | [15, 24] |
| Rhinopristiformes | Rhinobatidae | *Pseudobatos productus* | No | No | 185.0 | 0.095 | 7.7 | 24.3 | 16.0 | 3.0 | 1 | 3.62 | 0.230 | [15, 25-27] |
| Rhinopristiformes | Rhinobatidae | *Rhinobatos rhinobatos* | No | Yes | 185.0 | 0.222 | 3.2 | 16.0 | 12.0 | 3.8 | 1 | 2.00 | 0.537 | [15, 28-33] |
| Rhinopristiformes | Trygonorrhinidae | *Zapteryx brevirostris* | No | No | 66.0 | 0.120 | 9.6 | 15.2 | 12.4 | 2.0 | 1 | 1.50 | 0.155 | [1] |
| Rhinopristiformes | Trygonorrhinidae | *Zapteryx exasperata* | No | No | 103.0 | 0.159 | 7.5 | 19.8 | 13.7 | 4.0 | 1 | 3.10 | 0.261 | [1] |
| Squaliformes | Centrophoridae | *Centrophorus granulosus* | No | No | 128.0 | 0.134 | 12.2 | 25.0 | 18.6 | 1.0 | 2 | 1.83 | 0.035 | [1] |
| Squaliformes | Centrophoridae | *Centrophorus squamosus* | No | No | 145.0 | 0.025 | 44.0 | 70.0 | 57.0 | 8.1 | 2.5 | 1.63 | 0.054 | [1] |
| Squaliformes | Centrophoridae | *Centroselachus crepidater* | No | No | 99.5 | 0.072 | 20.0 | 57.0 | 38.5 | 6.0 | 2 | 1.62 | 0.101 | [1] |
| Squaliformes | Centrophoridae | *Deania calcea* | No | No | 119.0 | 0.077 | 25.0 | 35.0 | 30.0 | 13.0 | 4 | 0.75 | 0.077 | [1] |
| Squaliformes | Dalatiidae | *Dalatias licha* | No | No | 182.0 | 0.198 | 6.8 | 18.4 | 12.6 | 12.0 | 3 | 0.90 | 0.221 | [1] |
| Squaliformes | Etmopteridae | *Etmopterus baxteri* | No | No | 88.0 | 0.038 | 30.0 | 57.0 | 43.5 | 11.0 | 3 | 1.50 | 0.076 | [1] |
| Squaliformes | Etmopteridae | *Etmopterus spinax* | No | No | 46.0 | 0.220 | 5.0 | 7.0 | 6.0 | 6.8 | 2 | 2.00 | 0.183 | [1] |
| Squaliformes | Squalidae | *Squalus acanthias* | No | Yes | 110.0 | 0.089 | 12.5 | 30.0 | 21.3 | 8.0 | 2 | 0.25 | 0.147 | [1] |
| Squaliformes | Squalidae | *Squalus blainvillei* | No | No | 92.0 | 0.102 | 5.1 | 15.0 | 10.1 | 4.0 | 2 | 1.00 | 0.178 | [1] |
| Squaliformes | Squalidae | *Squalus megalops* | No | No | 78.2 | 0.032 | 15.0 | 32.0 | 23.5 | 3.0 | 2 | 1.70 | 0.081 | [1] |
| Squaliformes | Squalidae | *Squalus mitsukurii* | No | No | 91.0 | 0.041 | 15.0 | 27.0 | 21.0 | 3.6 | 2 | 2.00 | 0.085 | [1] |
| Squatiniformes | Squatinidae | *Squatina californica* | No | No | 118.0 | 0.162 | 10.0 | 35.0 | 22.5 | 6.0 | 1 | 0.75 | 0.214 | [1] |
| Squatiniformes | Squatinidae | *Squatina dumeril* | No | No | 152.0 | 0.015 | 25.0 | 34.4 | 29.7 | 4.0 | 1 | 0.92 | 0.104 | [1] |
| Squatiniformes | Squatinidae | *Squatina guggenheim* | No | No | 92.0 | 0.275 | 4.0 | 12.0 | 8.0 | 5.5 | 3 | 2.00 | 0.184 | [1] |
| Squatiniformes | Squatinidae | *Squatina occulta* | No | No | 131.0 | 0.129 | 10.0 | 21.0 | 15.5 | 6.0 | 4 | 3.00 | 0.097 | [1] |
| Torpediniformes | Torpedinidae | *Torpedo californica* | No | No | 102.0 | 0.073 | 9.0 | 16.0 | 12.5 | 17.0 | 1 | 1.83 | 0.289 | [1] |
| Torpediniformes | Torpedinidae | *Torpedo marmorata* | No | No | 63.0 | 0.088 | 12.5 | 20.0 | 16.3 | 11.0 | 3 | 1.70 | 0.128 | [1] |
| Torpediniformes | Torpedinidae | *Torpedo torpedo* | No | No | 41.0 | 0.275 | 4.0 | 10.0 | 7.0 | 3.4 | 1 | 8.50 | 0.264 | [1] |
