## Supplementary figures and images for "Population productivity of wedgefishes, guitarfishes, and banjo rays: inferring the potential for recovery"

### S2 Figure

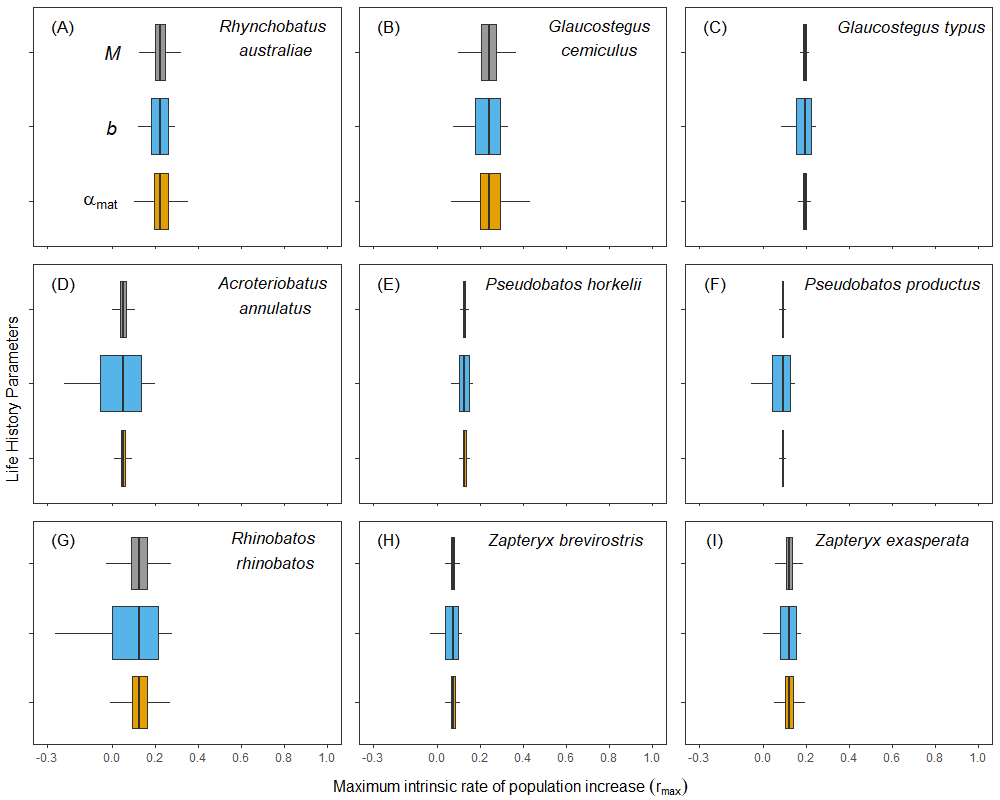

### S3 Figure

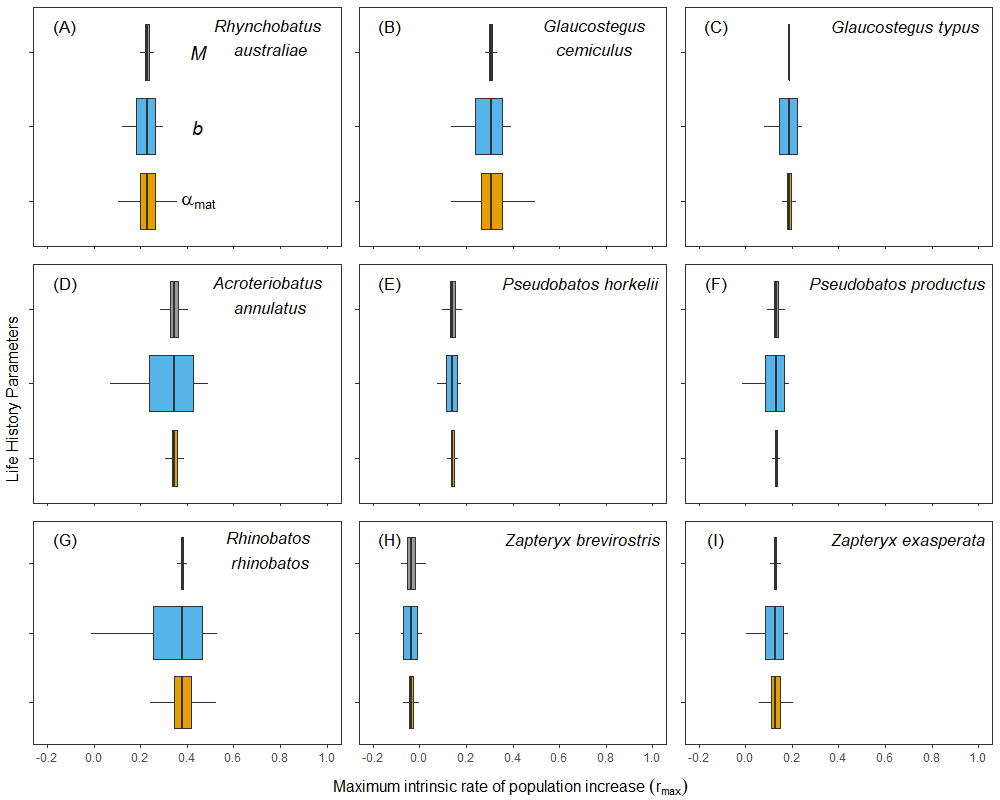

### S4 Figure

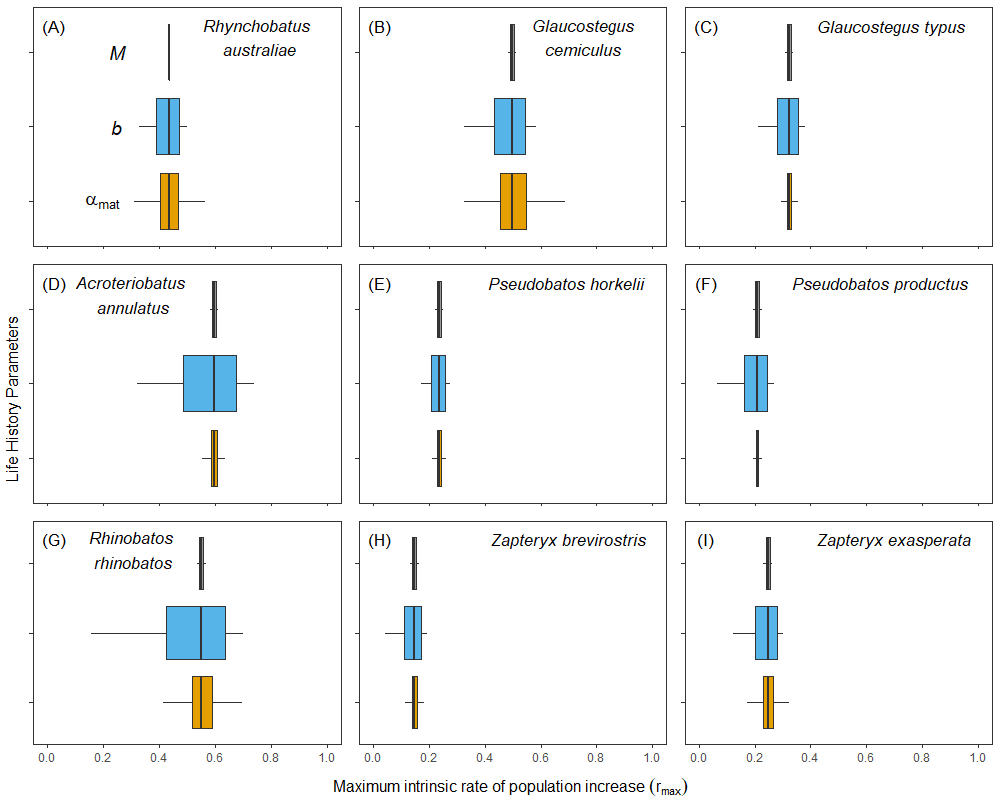
